# Distinct DAXX effector modules separate H3.3 nucleosome assembly from ERV silencing

**DOI:** 10.64898/2026.03.21.713205

**Authors:** Aayushi Y. Jain, Dominic Hoelper, Andrew Q. Rashoff, Peter W. Lewis

**Affiliations:** Department of Biomolecular Chemistry, School of Medicine and Public Health, University of Wisconsin, Madison, WI, 53706

## Abstract

Endogenous retroviruses (ERVs) compromise genome integrity when expressed, and cells have evolved chromatin-based pathways to silence their transcription. The histone H3.3 chaperone DAXX localizes to a subset of ERVs and enforces their silencing through incompletely defined mechanisms. Using complementary biochemical and genetic approaches, we identify a conserved basic patch within the DAXX histone-binding domain that engages DNA, promotes H3.3 nucleosome assembly in vitro, and is required for H3.3 enrichment at DAXX-bound ERVs in cells. Despite failure to deposit H3.3, DAXX with substitutions in this basic patch retains localization to ERVs and preserves silencing, indicating that histone H3.3 is dispensable for DAXX-mediated repression of ERVs. By contrast, ERV silencing requires the DAXX C-terminal SUMO-interacting motif, which mediates recruitment of SUMOylated repressors, including MORC3. These findings define modular outputs downstream of DAXX recruitment that uncouple nucleosome assembly from ERV silencing and highlight SUMO-dependent effector recruitment as the primary mechanism of silencing.

## INTRODUCTION

Silent chromatin states comprise heterochromatin domains that limit engagement of the transcriptional machinery and reinforce genome organization^1,2^. In mammalian cells, H3K9me3 together with conserved effector proteins forms a central axis for heterochromatin assembly and maintenance across repetitive DNA, including satellite and tandem repeats, and transposable element insertions^3,4^. Dynamic regulation of H3K9me3-marked heterochromatin at genes and repeats contributes to lineage specification and maintenance of cell identity^5–7^, in part by constraining repetitive DNA that is intrinsically prone to transcriptional activation and recombination^2,8–10^.

Among transposable elements, endogenous retroviruses (ERVs) are a prominent long terminal repeat class whose sequences can function as promoters or enhancers when chromatin repression is relaxed^11,12^. ERV silencing both limits retroelement transcription and insulates nearby gene expression programs from *cis*-regulatory activities embedded within ERV insertions^13–15^. This restraint is implemented by layered chromatin-mediated mechanisms, including H3K9me3-based heterochromatin, histone deacetylation, and complementary silencing systems that preferentially act at recently integrated, lineage-restricted, and low-divergence retroelements^13,16,17^. Beyond these enzymatic and reader-mediated pathways, prior work suggests that nucleosome composition at ERVs, including incorporation of histone variants, contributes to silencing, but the biochemical mechanism remains unresolved^18,19^.

Replication-independent nucleosome replacement, including histone-variant exchange, offers a route to modulate chromatin structure at ERVs and other repeats outside S phase^20–26^. Histone H3.3 is a replacement H3 deposited through at least two major chaperone pathways and is enriched at dynamic euchromatin as well as select heterochromatic and repetitive domains, including telomeres^27–34^. In embryonic stem cells (ESCs), H3.3-containing nucleosomes are enriched at a subset of ERVs within H3K9me3-marked chromatin, and prior studies have found that H3.3 deposition pathways are necessary for silencing specific ERV families^35–38^. However, whether H3.3 nucleosome assembly is required for ERV silencing, rather than reflecting a dynamic feature of the underlying silent chromatin environment, remains incompletely defined.

The function of H3.3 in repeat silencing can be interrogated through the DAXX H3.3-deposition pathway, which targets H3.3 to heterochromatic, repeat-rich loci. DAXX is an H3.3-selective histone chaperone that often functions with ATRX to support replication-independent H3.3 nucleosome assembly at these regions^28–31,39–43^. Accordingly, the ATRX-DAXX complex is widely viewed as a major route for H3.3 incorporation at repetitive DNA. Consistent with this role, prior studies implicate ATRX and DAXX in silencing specific murine ERV families, linking H3.3 deposition pathways to retroelement control^35–38,44–47^. The importance of defining DAXX function at repetitive chromatin is further underscored by recurrent *DAXX* alterations in tumors associated with genome instability and aberrant telomere maintenance^48–54^. However, DAXX has functions beyond H3.3 deposition, including SUMO-dependent interactions with chromatin repressors implicated in ERV silencing^55^. Notably, genetic ablation of H3.3 in mammalian cells markedly reduces DAXX steady-state abundance, an effect reversed by H3.3 re-expression^37^. This coupling complicates interpretation of H3.3 knockout phenotypes at ERVs, because loss of H3.3 perturbs both nucleosome composition and DAXX abundance, making it difficult to distinguish a direct requirement for H3.3 nucleosome assembly from mechanistically distinct DAXX-dependent repression functions.

Here, we define the genomic targeting landscape of DAXX in mouse ESCs (mESCs) and delineate distinct molecular determinants that govern H3.3 deposition versus ERV silencing. Genome-wide chromatin profiling shows that DAXX occupancy is concentrated at a subset of ERVK elements, enriched for evolutionarily young insertions. Using structure-guided mutagenesis and biochemical reconstitution, we identify a conserved basic surface within the DAXX histone-binding domain that mediates DAXX-DNA engagement. This surface is required for DAXX-dependent nucleosome assembly *in vitro* and for H3.3 enrichment at DAXX-bound ERVs in cells. At these ERVs, H3.3 enrichment occurs independently of ATRX, supporting an ERV-associated DAXX deposition pathway distinct from ATRX-dependent deposition. By contrast, ERV silencing depends on the DAXX C-terminal SUMO-interacting motif (SIM), which engages SUMOylated repressors and promotes MORC3 recruitment at DAXX-bound ERVs. By identifying and leveraging DAXX separation-of-function alleles, we show that H3.3 deposition is dispensable for ERV silencing, establishing that DAXX-directed nucleosome assembly and transcriptional repression are mechanistically separable post-recruitment outputs.

## RESULTS

### DAXX is enriched at young ERVK elements and is required for their repression in mESCs

To define the DAXX targeting landscape in mESCs and relate occupancy to local chromatin context and transcriptional output, we performed DAXX ChIP-seq with repeat-aware analyses, integrated these maps with chromatin profiles at bound loci, and compared RNA-seq outputs in *Daxx* knockout and wild-type rescue settings. High-confidence DAXX peaks mapped predominantly to repetitive DNA, with transposable elements comprising the largest annotated fraction and most remaining peaks distributed across intronic and intergenic regions (**Fig. 1A**). Repeat-family quantification showed that DAXX enrichment was concentrated within a restricted subset of repeat families rather than broadly distributed across repetitive elements genome-wide (**Fig. 1B**). Enrichment testing further supported preferential association of DAXX peaks with LTR elements (**Fig. S1A**), and within LTR retrotransposons, DAXX peaks were biased toward ERVK relative to other LTR classes (**Fig. S1B**). At ERVK subfamily resolution, DAXX occupancy concentrated on a limited set of lineages, with IAPEz-int and additional subclasses, including ETnERV2-int, representing prominent targets (**Fig. S1C**). Consistent with selective targeting, DAXX-bound ERVs were enriched for younger insertions compared with the genome-wide ERV distribution (**Fig. 1C**). These observations defined the ERVK loci used to test how DAXX chromatin occupancy is coupled to H3.3 enrichment and repression.

**Figure 1.**
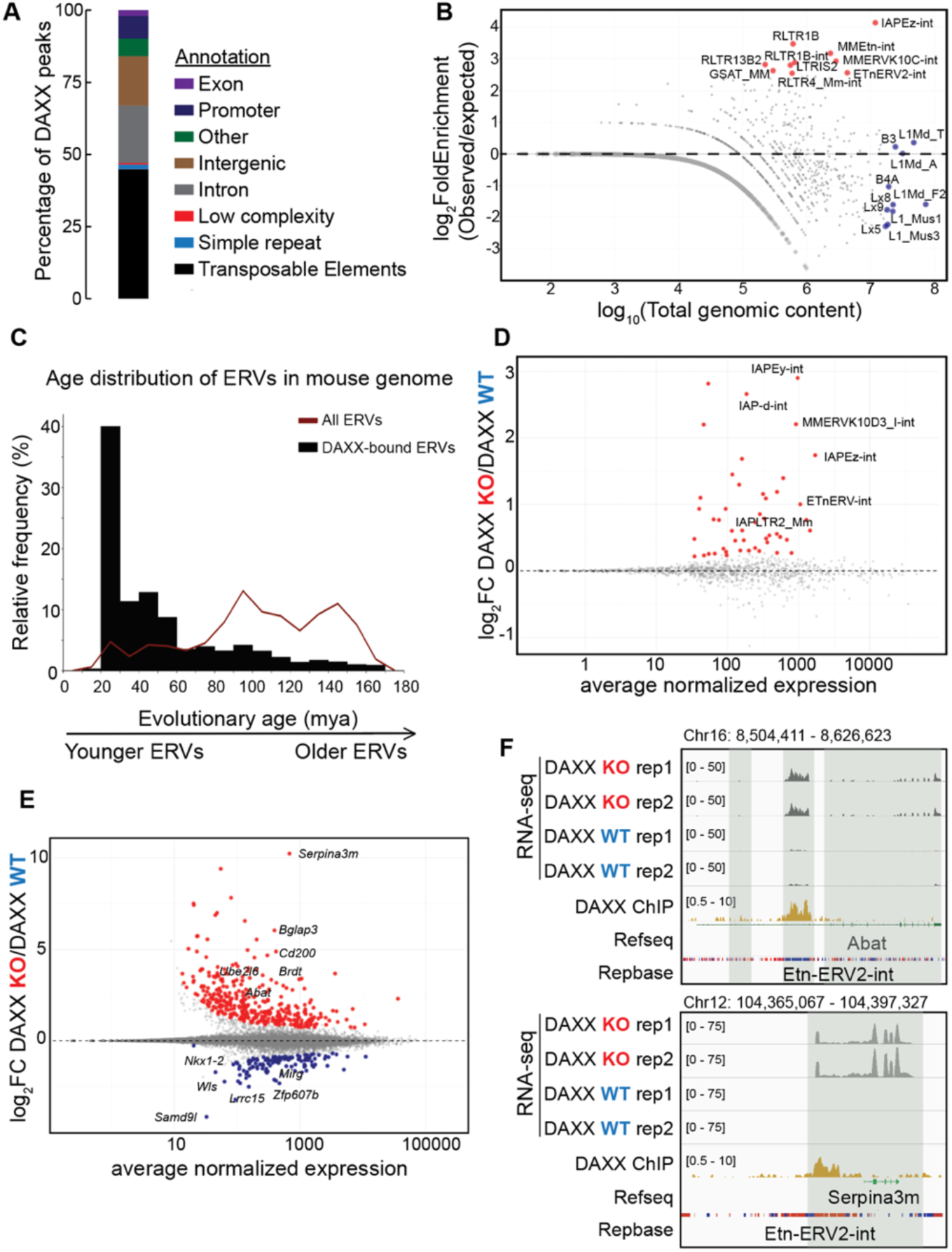
DAXX selectively localizes to young ERVK elements in mESCs and marks focal H3.3 enrichment at ERV loci. (**A**) Feature and repeat annotation of DAXX ChIP-seq peaks. (**B**) Repeat-family enrichment among DAXX peaks (log_2_ observed/expected versus genomic content; dashed line, observed/expected = 1; red, top DAXX-enriched repeats; blue, most abundant repeats in mouse genome). (**C**) Insertion age distributions for DAXX-bound versus all ERVs. (**D,E**) RNA-seq MA plots comparing *Daxx* knockout (KO) and wild-type (WT) rescue mESCs (n = 2 per condition) at the ERV family (**D**) and gene (**E**) levels (DESeq2; adjusted p < 0.05). (**F**) Genome browser views of EtnERV2-int insertions within *Abat* and *Serpina3m* showing DAXX occupancy and increased RNA-seq signal in KO; tracks include RNA-seq replicates, DAXX ChIP-seq, gene models, and RepeatMasker annotations.

We next asked whether DAXX-enriched repeats reside within chromatin contexts characteristic of established ERVK silencing paradigms in mESCs, which feature KAP1 and SETDB1 recruitment, co-occupancy by MORC3, SMARCAD1, and ATRX, and enrichment of H3K9me3 and H4K20me3 with comparatively low H3K27me3^13,35,45,46,56–59^. Peak-centered alignment of ChIP-seq profiles at DAXX peaks overlapping IAPEz-int insertions recapitulated this signature (**Fig. S1E**). Together, these data place DAXX-bound IAPEz-int elements within a heterochromatin-associated repressor landscape characteristic of transcriptionally silent ERVK loci in mESCs. To link DAXX occupancy with genome-wide H3.3 enrichment and transcriptional output, we integrated H3.3 ChIP-seq and RNA-seq using repeat-aware analyses anchored to the DAXX ChIP-seq maps. At DAXX peaks, H3.3 ChIP-seq signal was lost in *Daxx*-knockout cells and restored by the wild-type transgene (**Fig. S1F**). Differential expression analysis showed that loss of DAXX results in statistically significant upregulation of a defined subset of ERV families (**Fig. 1D**).

To evaluate whether gene-level transcriptional changes were enriched near DAXX binding sites, we stratified differentially expressed genes by distance to the nearest DAXX peak and found that approximately 80% of these genes lay within 100 kb of a DAXX peak (**Fig. 1E, 1F, S1D**). This enrichment is consistent with local *cis* effects of DAXX-bound ERV elements on transcription at nearby loci. Together, these data establish that DAXX is preferentially concentrated at young ERVK elements in mESCs, where it restrains both ERV expression and ERV-associated misregulation of neighboring genes.

### A conserved basic surface in DAXX Histone-binding domain promotes DNA engagement

Having established that DAXX localizes to young ERVK loci, where it represses repeat transcription and thereby preserves appropriate expression of ERV-proximal genes, we next asked how this site-specific occupancy is coupled to H3.3 enrichment and transcriptional silencing. Prior genetic and biochemical studies at ERVs left unresolved whether DAXX-dependent H3.3 enrichment is mechanistically linked to silencing or instead reflects a distinct post-recruitment activity. Structural studies established that the DAXX histone-binding domain (HBD) wraps an H3.3-H4 heterodimer, occluding extensive histone surface area and stabilizing a soluble pre-deposition complex^60–62^. However, productive nucleosome assembly requires this shielded intermediate to transition into a DNA-engaged state that supports (H3.3-H4) tetramer formation and histone transfer onto DNA^60–63^. How DAXX prevents premature histone-DNA association yet subsequently promotes DNA engagement and histone release during deposition, has remained unresolved. We therefore asked whether DAXX contains a separable DNA engagement surface that facilitates this handoff. This model is supported by precedent from the H3-H4 chaperone CAF-1, which harbors DNA interaction surfaces that facilitate deposition during nucleosome assembly^64–69^.

To identify candidate DNA-contacting regions, we started from the crystal structure of the DAXX HBD-H3.3-H4 ternary complex and used AlphaFold to model its engagement with Widom 601 DNA. In this configuration, the DNA duplex localized adjacent to a prominent basic region on the HBD, suggesting that this surface may participate in substrate DNA engagement (**Fig. 2A,B**). Electrostatic mapping of HBD within the ternary complex identified basic surface patches that are spatially separable from the histone-binding interface. These analyses converged on the same basic region, motivating a graded alanine-substitution series to neutralize this surface (BPM_4A and BPM_7A) (**Fig. 2C**). The substituted residues cluster within conserved segments of the HBD, consistent with the idea that this basic region represents a functionally important feature (**Fig. S2A,B**).

**Figure 2.**
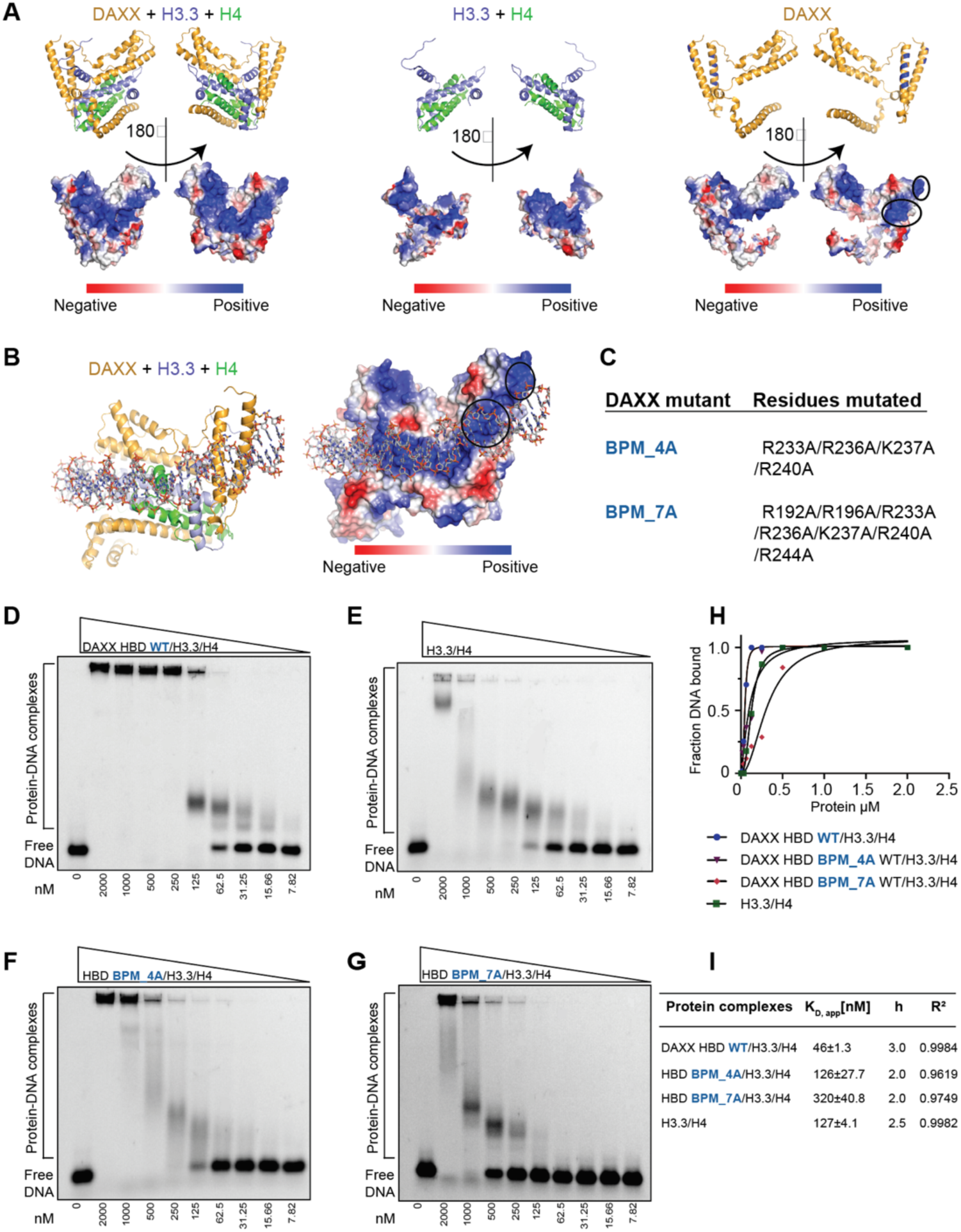
A basic surface on the DAXX histone-binding domain supports DNA engagement by the DAXX HBD–H3.3–H4 complex. (**A**) Crystal structure of DAXX HBD bound to H3.3–H4 (PDB: 4H9N) shown as ribbons and electrostatic surfaces in two orientations, highlighting basic patches on DAXX HBD that are distinct from the H3.3–H4 binding interface. (**B**) AlphaFold model of the DAXX HBD–H3.3–H4 complex positioned on Widom 601 DNA, with basic regions proximal to the modeled DNA trajectory. (**C**) Basic patch substitution series: BPM_4A (R233A, R236A, K237A, R240A) and BPM_7A (BPM_4A plus R192A, R196A, R244A). (**D–G**) EMSAs with Cy5-labeled Widom 601 DNA across a 7.8 nM to 2.0 μM titration, showing enhanced DNA shifting by DAXX HBD–H3.3–H4 relative to H3.3–H4 and reduced protein-DNA complex formation by BPM mutants, strongest effect for BPM_7A. (**H**) Bound DNA fraction quantified from three independent EMSAs and fit with a Hill model. (**I**) Apparent K_D_ and Hill coefficients from Hill fits (mean ± error from fitting).

We tested whether the HBD confers DNA binding to the HBD-H3.3-H4 pre-deposition complex using electrophoretic mobility shift assays (EMSAs) with Cy5-labeled Widom 601 DNA across a protein titration series. Because the HBD alone is insufficient to support tetrasome assembly^60–62^, shifted species under these conditions are most consistent with direct binding of the pre-deposition complex to DNA. To evaluate whether the shift depends on stable tetramer assembly, we repeated the same EMSAs using H3.3(L126A,I130A), a incorporation-defective mutant that disrupts the H3.3-H3.3 dimerization interface required for stable (H3.3-H4) tetramer formation and chromatin incorporation while preserving binding to DAXX (**Fig. S2C,D**)^37^. H3.3-H4 alone shifted DNA only at high protein concentrations, consistent with nonspecific histone-DNA association *in vitro* rather than productive tetrasome assembly under these conditions (**Fig. 2D-G**).

Relative to H3.3-H4 alone, addition of the HBD increased DNA shifting at lower protein concentrations, consistent with higher apparent DNA affinity of the HBD-H3.3-H4 pre-deposition complex (**Fig. 2D-E**). Neutralization of the basic surface reduced complex formation across the titration series (**Fig. 2D-G**). Notably, the HBD enhanced shifting to a similar extent with H3.3(L126A,I130A) (**Fig. S2C,D**). Because this mutant cannot support stable (H3.3-H4) tetramerization, the persistence of the shifted species indicates that the EMSA signal reflects direct DNA engagement by the DAXX HBD-H3.3-H4 pre-deposition complex rather than formation of a tetrasome or nucleosome-like particle.

We quantified these titrations by fitting binding data with a Hill-type model to capture the concentration dependence of complex formation (**Fig. 2H**), enabling estimation of apparent dissociation constants and Hill coefficients (**Fig. 2I**). The Hill coefficients of the DAXX HBD-H3.3-H4 complex indicate apparent cooperativity in DNA binding and/or higher-order complex formation. Similar behavior has been previously reported for CAF-1-H3-H4 on DNA substrates, where it was interpreted as coupled association of two chaperone-histone complexes on a single DNA molecule during nucleosome assembly^64,66–68^. Neutralization of the basic surface caused graded rightward shifts in the binding curves and reduced apparent affinity, consistent with impaired DNA engagement by mutant ternary complexes (**Fig. 2H,I**).

Together, these data identify a conserved basic surface on DAXX that enhances DNA engagement by the HBD-H3.3-H4 complex *in vitro*. This result provides a mechanistic basis to test whether the same surface contributes to nucleosome assembly and to restoration of H3.3 enrichment at DAXX-bound loci in cells.

### DAXX DNA engagement promotes H3.3 nucleosome assembly *in vitro* and enrichment *in vivo*

Because the DAXX basic patch binds DNA, we next asked whether it is also required for H3.3-H4 deposition onto DNA for nucleosome assembly. We and others have previously used plasmid supercoiling and related DNA topology assays to assess DAXX-dependent tetrasome assembly *in vitro*^28,29^, in which tetramer assembly generates negative supercoils that are recovered as distinct topoisomers after relaxation and deproteinization. Increasing amounts of full-length wild-type DAXX-H3.3-H4 produced a progressive increase in negatively supercoiled plasmid species, consistent with dose-dependent nucleosome assembly. By contrast, complexes harboring basic patch substitutions displayed reduced supercoiling across the same titration series (**Fig. 3A**, **S3C**). We tested both 4A and 7A basic patch substitution series *in vitro*, and for all subsequent cell-based experiments we used the 4A allele, hereafter termed the basic patch mutant (BPM). Together, these results indicate that the conserved basic surface is essential for productive nucleosome assembly *in vitro*, consistent with a role in facilitating engagement of DAXX-H3.3-H4 with DNA during deposition.

**Figure 3.**
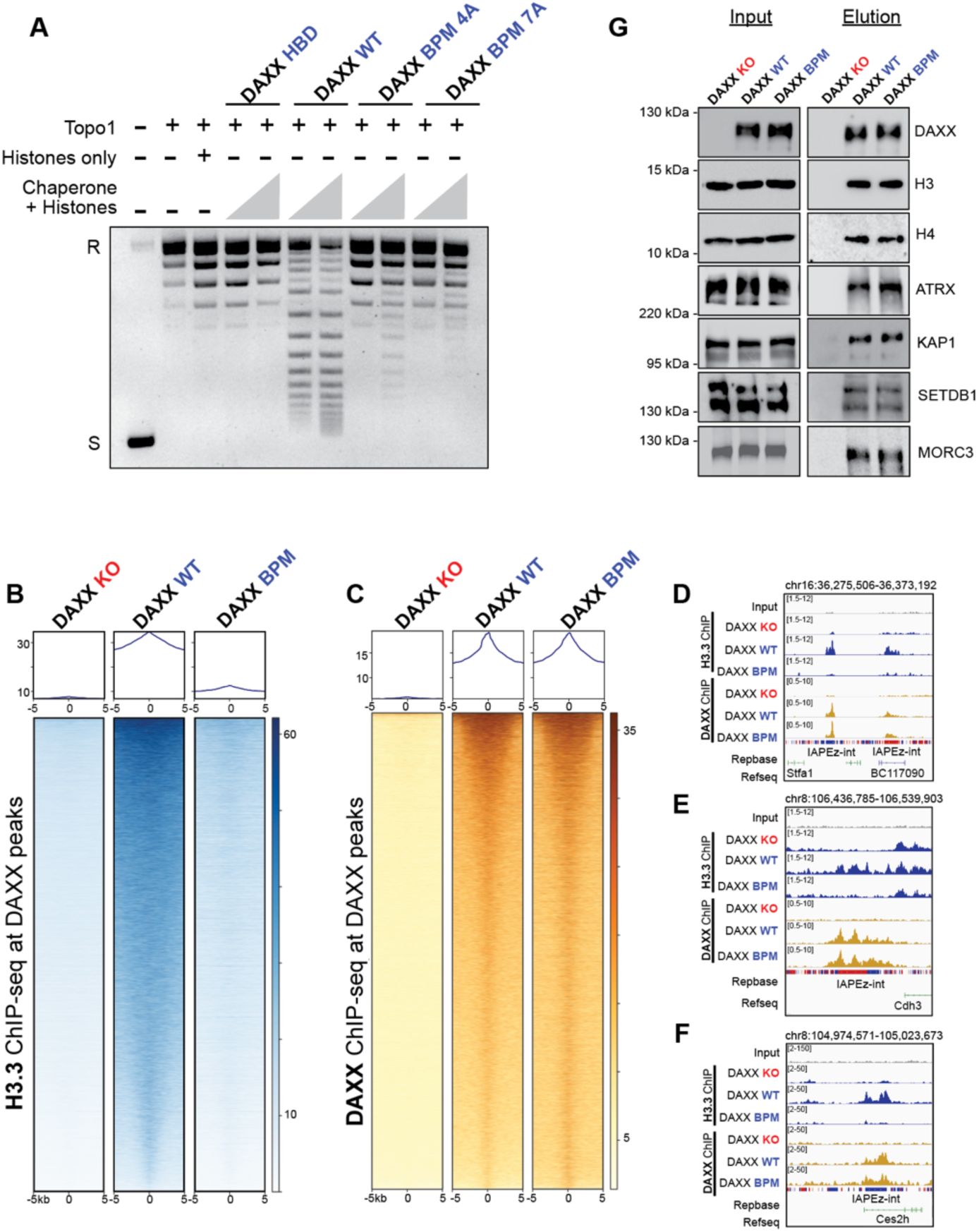
A basic surface within the DAXX histone-binding domain promotes nucleosome assembly *in vitro* and H3.3 enrichment in cells. (**A**) Plasmid supercoiling assay after Topo1 relaxation. Increasing amounts of DAXX–H3.3–H4 complexes generate a topoisomer ladder consistent with dose-dependent nucleosome assembly, whereas basic patch mutants (BPM_4A and BPM_7A; Fig. 2C) show reduced supercoiling, strongest effect for BPM_7A. (**B**) H3.3 ChIP-seq metaprofiles and heat maps centered on DAXX peak summits (±5 kb), showing reduced H3.3 in *Daxx* KO cells, restoration by wild-type rescue, and impaired restoration by the basic patch mutant (BPM). (**C**) DAXX ChIP-seq at the regions in (**B**), showing loss in KO and restoration in both WT and BPM rescues. (**D–F**) Genome browser views at representative loci showing restored DAXX occupancy in WT and BPM rescues with reduced H3.3 signal in BPM relative to WT, alongside input, RepeatMasker, and RefSeq annotations. (**G**) Anti-FLAG immunoprecipitation from nuclear extracts of *Daxx* KO cells or cells expressing FLAG-DAXX WT or BPM, showing comparable co-purification of histone H3 and H4 and retained detection of ATRX, KAP1, SETDB1, and MORC3 in WT and BPM eluates.

We then tested whether the same basic surface is required for H3.3 enrichment at endogenous DAXX targets in mESCs. At DAXX-enriched peaks, H3.3 ChIP-seq signal was low in *Daxx*-knockout cells and restored by wild-type DAXX rescue, whereas rescue with the BPM failed to restore H3.3 enrichment (**Fig. 3B**). This deposition defect is not explained by impaired targeting, as DAXX ChIP-seq showed comparable occupancy of wild-type and BPM at these peaks (**Fig. 3C**). Consistent with a locus-restricted role, H3.3 profiles at DAXX-independent regions were similar across knockout and rescue conditions (**Fig. S3H**). Genome browser views recapitulated these trends by showing preserved DAXX occupancy with impaired H3.3 restoration at representative repeat-associated loci in the BPM rescue (**Fig. 3D-F**).

Because H3.3 deposition can be influenced by altered DAXX abundance, turnover, or complex integrity, we evaluated alternative explanations for the BPM deposition phenotype. Immunoblotting revealed comparable steady-state levels of wild-type and BPM transgenes, whereas a histone-binding mutant (HBM) accumulated at lower levels (**Fig. S3A**), consistent with prior evidence that histone engagement stabilizes DAXX in cells^37^. Cycloheximide chase assays showed similar decay kinetics for wild-type DAXX and BPM over a 24-hour time course, indicating comparable protein half-life (**Fig. S3B**). FLAG immunoprecipitation from nuclear extracts recovered histone H3 and histone H4 with both wild-type DAXX and BPM and retained detectable association with ATRX, KAP1, SETDB1, and MORC3 (**Fig. 3G**). To test whether the deposition defect reflects a general reduction in HBD basic charge or a site-specific requirement, we additionally defined a second, spatially distinct basic surface on the HBD (hereafter “AltBP”; **Fig. S3E**) and generated a six-alanine substitution series (AltBP_6A). In locus-level ChIP-qPCR, AltBP_6A behaved like wild-type, restoring H3.3 enrichment at representative DAXX-bound ERVs (**Fig. S3G**), in contrast to neutralization of the primary basic patch (BPM series), which produced a pronounced defect in H3.3 deposition. In parallel, RT-qPCR analysis of H3.3 deposition-dependent, DAXX-regulated transcriptional targets showed that BPM resembled the *Daxx*-knockout state at deposition-sensitive loci (**Fig. S3D-H**)^37^. Collectively, these controls argue against altered expression, reduced DAXX stability, loss of histone association, or broad disruption of partner interactions as primary explanations for the deposition defect and instead support a model in which the HBD basic patch mediates a post-recruitment step that enables productive H3.3-H4 transfer onto DNA by facilitating chaperone-DNA contact.

### Genetic separation uncouples ERV recruitment from H3.3 enrichment by DAXX

Building on the finding that the basic patch within the histone-binding domain is required for H3.3 enrichment, we next asked which additional DAXX regions distinguish chromatin recruitment from productive H3.3-H4 deposition at DAXX-enriched repeat loci. A central question was whether H3.3 enrichment at these ERVs depends on the canonical ATRX-DAXX partnership, since prior genetic and biochemical work had suggested that DAXX may act at ERVs through an ATRX-independent pathway^37^. To guide selection of candidate determinants, we integrated structural information with AlphaMissense scores^70^, which predict the functional impact of missense variation and highlight regions of elevated predicted deleteriousness across human DAXX. This analysis pointed to the N-terminal four-helix bundle, the HBD basic surface, and the C-terminal SIM, aligning with domains and interfaces tested experimentally here (**Fig. S8C**). We therefore expressed a panel of structure-informed DAXX point mutants and deletions in *Daxx*-knockout mESCs (**Fig. 4A; Fig. S6A–D**) and quantified H3.3 profiles at DAXX peaks at ERVs by ChIP-seq. Strikingly, a previously characterized ATRX-binding mutant DAXX transgene^37^ fully restored H3.3 enrichment at DAXX peaks (**Fig. 4B,C; Fig. S4A**), indicating that ATRX binding is not required for DAXX-dependent H3.3 accumulation at these ERVs. Consistent with this conclusion, a transgene initiating at the HBD, thereby deleting the entire N-terminal region, including 4HB (HBD-end), likewise restored the H3.3 signal in cells. Together, these results indicate that ATRX association and N-terminal regions to the HBD are dispensable for DAXX-mediated H3.3 accumulation at repeats in mESCs.

**Figure 4.**
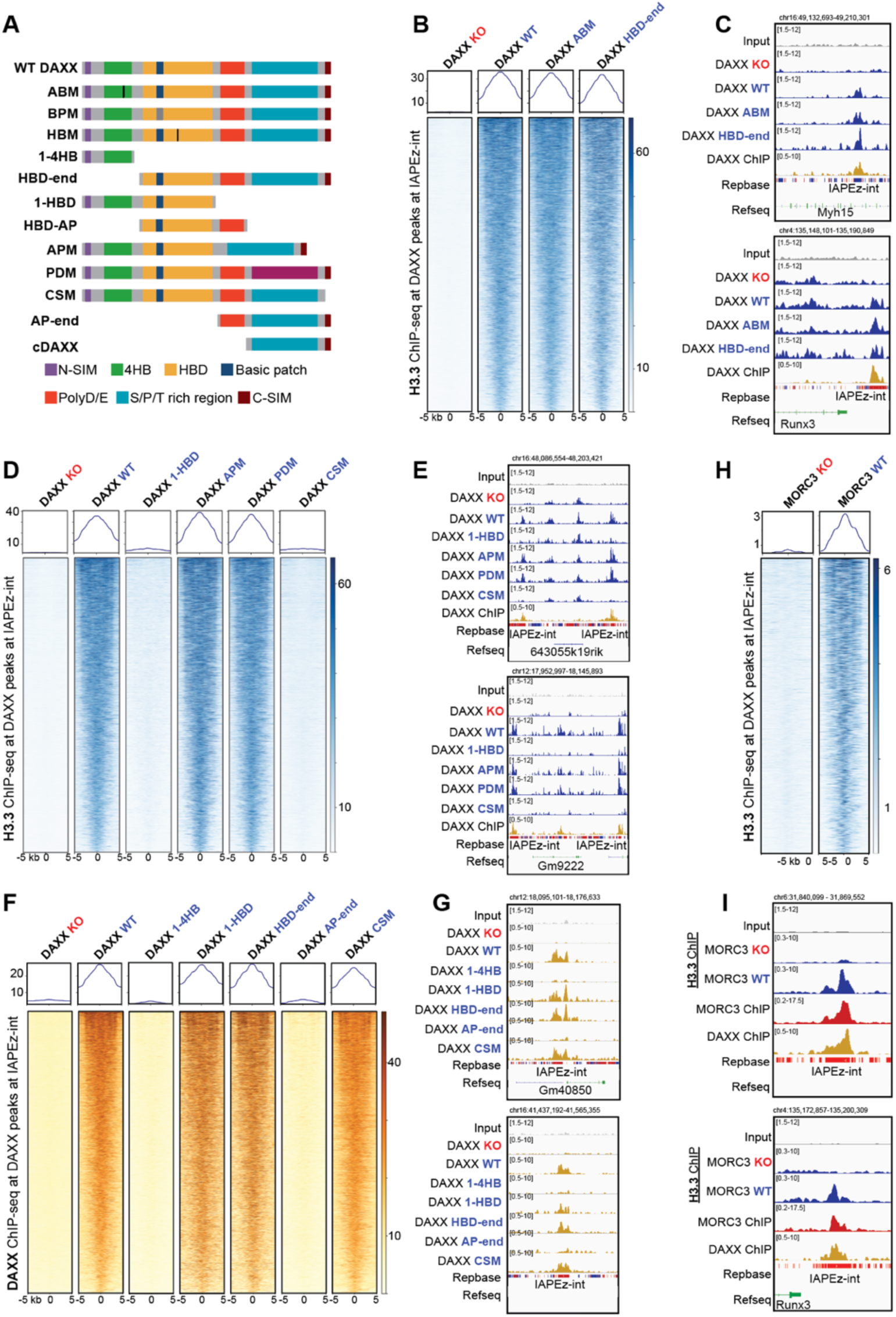
The C-terminal SIM and MORC3 are required for H3.3 enrichment, whereas the HBD is required for both recruitment of DAXX to ERVs and H3.3 deposition. (**A**) DAXX domain map and constructs panel, including N-SIM, four-helix bundle (4HB), HBD (with basic patch), polyD/E region, S/P/T-rich region, and C-SIM, with ABM, BPM, APM, PDM, CSM, and truncations as indicated. (**B**) H3.3 ChIP-seq signal at DAXX-bound IAPEz-int loci, showing loss in *Daxx*-knockout cells and restoration upon rescue with wild-type DAXX, ABM, or HBD-end. (**C**) Browser views of a DAXX-dependent IAPEz-int locus and a DAXX-independent control. (**D,E**) H3.3 ChIP-seq across additional constructs, showing that C-terminal sequences including the C-SIM are required for H3.3 restoration (1-HBD and CSM fail to rescue; APM and PDM rescue). (**F,G**) DAXX ChIP-seq showing HBD-dependent recruitment, with CSM retaining ERV occupancy. (**H,I**) H3.3 ChIP-seq and browser views showing reduced H3.3 at DAXX-bound ERVs in MORC3 KO mESCs.

Because the HBD alone is insufficient for H3.3 deposition *in vitro*, we then focused on sequences C-terminal to the HBD, including the polyD/E acidic tract and the disordered tail that contains a C-terminal SUMO-interacting motif (C-SIM). In prior work, we reported that the pancreatic neuroendocrine tumor specimen MK38 harbors a premature stop mutation in DAXX that removes the final eight amino acids and thereby deletes the C-SIM (**Fig. S8B**)^71^. Consistent with this functional importance, disruption of the motif eliminated punctate nuclear localization and instead produced a diffused nuclear signal for DAXX (**Fig. S8A**). The polyD/E region has been implicated in an ATP-independent chaperone-like activity of DAXX in protein folding and disaggregation, supporting the broader idea that the C-terminal half of DAXX can provide functional outputs distinct from histone binding by the HBD^72^. Full-length mutants targeting the acidic patch (APM) or the phosphorylation-site cluster (PDM) restored H3.3 ChIP-seq enrichment at DAXX peaks to levels comparable to wild-type rescue. By contrast, a truncation terminating near the acidic region (1-HBD) and the C-terminal SIM mutant (CSM) failed to restore H3.3 enrichment (**Fig. 4D,E**). These results suggest that the acidic patch integrity and the phosphosite cluster were dispensable, whereas sequences within the broader C-terminal region, including the C-SIM, were required for H3.3 enrichment at DAXX-enriched loci in cells.

To separate DAXX recruitment from H3.3-deposition defects, we measured DAXX occupancy across the same construct panel. DAXX ChIP-seq showed that localization to repeat-associated sites required the HBD: constructs lacking the HBD showed little to no enrichment, whereas HBD-containing constructs localized robustly (**Fig. 4F,G**). Notably, the CSM mutant retained DAXX occupancy but failed to restore H3.3 enrichment, indicating that the C-SIM functions downstream of chromatin recruitment rather than in controlling DAXX localization. These data support a stepwise model in which the HBD is required for targeting and stable occupancy at ERV-linked loci, whereas C-terminal determinants provide a distinct post-recruitment activity that is necessary to convert DAXX occupancy into local H3.3 enrichment. This interpretation aligns with prior structural and biochemical work showing that the isolated DAXX HBD stabilizes an H3.3-H4 pre-deposition complex but is insufficient, on its own, to drive efficient nucleosome assembly *in vitro*^60–62^. Our rescue-mapping experiments extend that framework in cells by demonstrating a requirement for C-terminal determinants beyond the HBD for H3.3 enrichment at endogenous targets.

### DAXX C-terminal SIM recruits MORC3 to DAXX-bound ERVs

Because loss of the C-SIM uncouples DAXX occupancy from local H3.3 enrichment (**Fig. 4F,G**), we asked whether the C-SIM engages SUMOylated effectors that act downstream of recruitment to promote H3.3 enrichment. Among candidate SUMOylated DAXX-interacting factors linked to ERV silencing, MORC3 is a SUMOylated ATPase implicated in repression of specific ERV families^46,73–75^. To examine the genomic relationship between MORC3 and DAXX directly, we performed MORC3 ChIP-seq. MORC3 co-enriched with DAXX at IAPEz-int peaks and, more broadly, overlapped with DAXX genome-wide, with MORC3 detected at nearly all DAXX peaks (**Fig. S1E; Fig. S7A-F**). Consistent with C-SIM-dependent engagement, MORC3 co-immunoprecipitated with wild-type DAXX but not with CSM^46^, whereas KAP1 and SETDB1 remained detectable in both conditions (**Fig. S4B**). Proteomic analyses similarly showed reduced recovery of SUMO1/2/3 and multiple SUMOylated proteins, including MORC3 and PML, with DAXX CSM (**Fig. S5D**). Together, these results indicate that the C-SIM supports association between DAXX and a subset of SUMOylated partners, including MORC3, without broadly disrupting interactions with other ERV-associated silencing factors.

To test whether DAXX recruits MORC3 at ERVs, we profiled MORC3 occupancy in *Daxx*-knockout mESCs and after rescue with wild-type DAXX or the CSM transgene. MORC3 ChIP-seq signal decreased in *Daxx*-knockout cells and was restored by wild-type rescue, but not by CSM (**Fig. 5E**). Because DAXX occupancy at ERVs was preserved in the CSM transgene (**Fig. 4F,G**), these data place MORC3 recruitment downstream of DAXX chromatin binding and identify the C-SIM as required for MORC3 localization at DAXX-bound ERVs (**Fig. 4F,G**; **Fig. 5E; Fig. S5E,F**).

**Figure 5.**
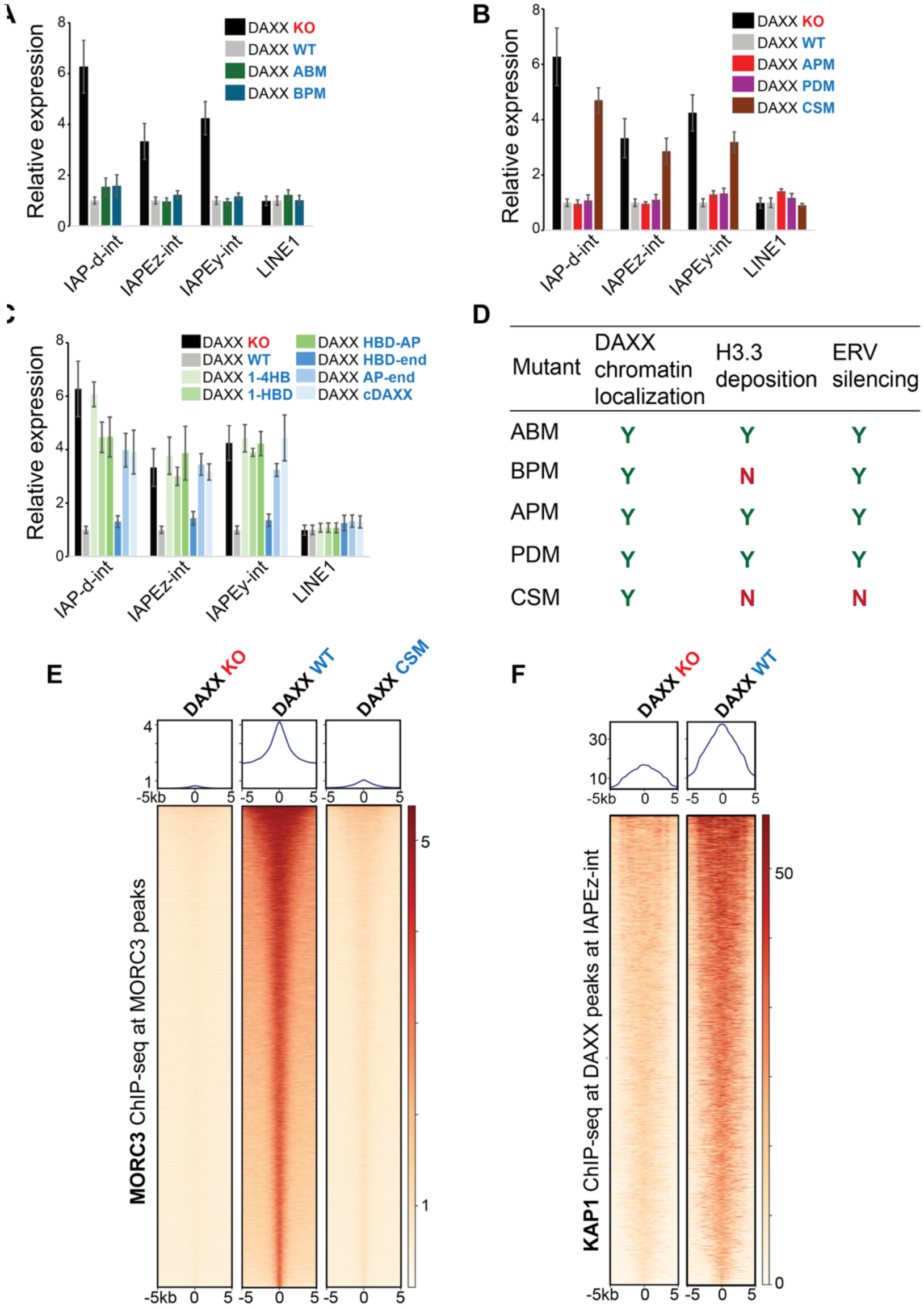
ERV silencing and MORC3 occupancy at ERVs both depend on the DAXX C-terminal SIM. (**A**-**C**) RT-qPCR of ERV and ERV-linked transcripts in *Daxx*-knockout and rescue mESCs across mutant series. ABM and BPM repress IAP-d-int, IAPEz-int, and IAPEy-int similarly to wildtype, whereas CSM and truncations lacking C-terminal function fail to repress multiple targets and resemble knockout cells. Constructs lacking the HBD show limited silencing. LINE-1 changes are modest. (**D**) Summary matrix of chromatin localization, H3.3 enrichment, and ERV silencing across the construct panel, highlighting recruitment-defective, enrichment-defective but silencing-competent, and recruitment-competent but silencing-defective states. (**E**) MORC3 ChIP-seq metaprofiles and heat maps showing reduced MORC3 occupancy in *Daxx*-knockout cells, restoration by wildtype rescue, and failure of CSM to restore MORC3 binding. (**F**) KAP1 ChIP-seq at DAXX-enriched IAPEz-int loci showing reduced occupancy in *Daxx*-knockout cells and restoration upon wildtype rescue.

We next tested whether MORC3 contributes to H3.3 enrichment at DAXX-bound ERVs. In *Morc3*-mESCs, H3.3 ChIP-seq signal was reduced at IAPEz-int-associated DAXX peaks, with little change at DAXX-independent loci (**Fig. 4H,I; S5D**). Wild-type DAXX restored H3.3 enrichment at DAXX peaks, whereas the CSM transgene did not, despite comparable DAXX occupancy (**Fig. S5A; Fig. 4F,G**). Together, these data support a post-recruitment, C-SIM-dependent step in which MORC3 engagement couples DAXX occupancy to local H3.3 enrichment at ERVs (**Fig. S5A-C; Fig. 5E**).

### ERV silencing requires DAXX C-terminal SIM-dependent effector engagement

We next asked whether DAXX C-terminal SIM-dependent effector engagement, inferred in part from MORC3 recruitment, is required for silencing at DAXX-bound ERVs. In transgene complementation assays, the ATRX-binding mutant (ABM) and the basic patch mutant (BPM) maintained silencing of ERVs and ERV-linked differentially expressed genes **(Fig. 5A, S6E**). This aligns with prior evidence that incorporation-defective H3.3 (L126A,I130A) preserves DAXX-dependent ERV silencing^37^. By contrast, CSM failed to restore silencing (**Fig. 5B**), and truncation analysis similarly mapped silencing activity to SIM-competent constructs, with HBD-end showing the strongest restoration among the truncations tested (**Fig. 5C**). Together, these data indicate that reduced H3.3 enrichment is not sufficient to abolish repression and instead identify a recruitment-competent, SIM-dependent effector arm as the major determinant of silencing output (**Fig. 5D**). Together with the recruitment and H3.3 enrichment data above, these experiments resolve three separable DAXX outputs at ERVs: HBD-dependent chromatin occupancy, basic patch-dependent H3.3 deposition, and C-SIM-dependent effector recruitment required for repression.

Consistent with a SIM-dependent effector mechanism, MORC3 occupancy at DAXX-bound ERVs required DAXX and the C-terminal SUMO-interacting motif (**Fig. 5E**). These findings support a model in which SIM-dependent engagement of SUMOylated effectors, exemplified by MORC3, constitutes a major repression arm at these loci that is distinct from DAXX-directed H3.3 deposition. Separately, KAP1 occupancy at DAXX-enriched IAPEz-int loci decreased in *Daxx*-knockout cells and increased upon wild-type DAXX reintroduction (**Fig. 5F**), consistent with reciprocal reinforcement between DAXX and KAP1 after localization at ERVs. Prior work reported that H3.3 loss reduces nuclear DAXX abundance and diminishes recruitment of both DAXX and KAP1 to ERVs^35,37^, raising the possibility that reduced KAP1 occupancy in H3.3-null cells may partly reflects reduced DAXX stability. We therefore favor a positive feedback model in which DAXX-enriched chromatin promotes KAP1 occupancy, and KAP1-associated repression machinery in turn sustains DAXX pathway activity, thereby mutually reinforcing a local repressive state. Together, these data place the DAXX C-terminal SIM downstream of chromatin occupancy in the repression pathway at DAXX-bound ERVs.

## DISCUSSION

ERV silencing in pluripotent cells is often framed as a coupled process in which repeat targeting, histone-variant deposition, and effector recruitment are mutually reinforcing. Here we demonstrate that, at DAXX-bound ERVs in mESCs, these outputs can be genetically separated. A conserved basic patch within the DAXX histone-binding domain promotes DNA engagement and is required for productive H3.3 nucleosome assembly and local H3.3 enrichment, yet disruption of this module preserves steady-state repression of ERVs. By contrast, silencing requires the DAXX C-terminal SUMO-interacting motif, which acts downstream of chromatin occupancy to support recruitment of MORC3 and likely additional SUMOylated effectors, whereas ATRX association is dispensable at these loci. Together, these findings support a model in which DAXX occupancy initiates separable post-recruitment pathways for H3.3 deposition and transcriptional silencing (**Fig. 6**).

**Figure 6.**
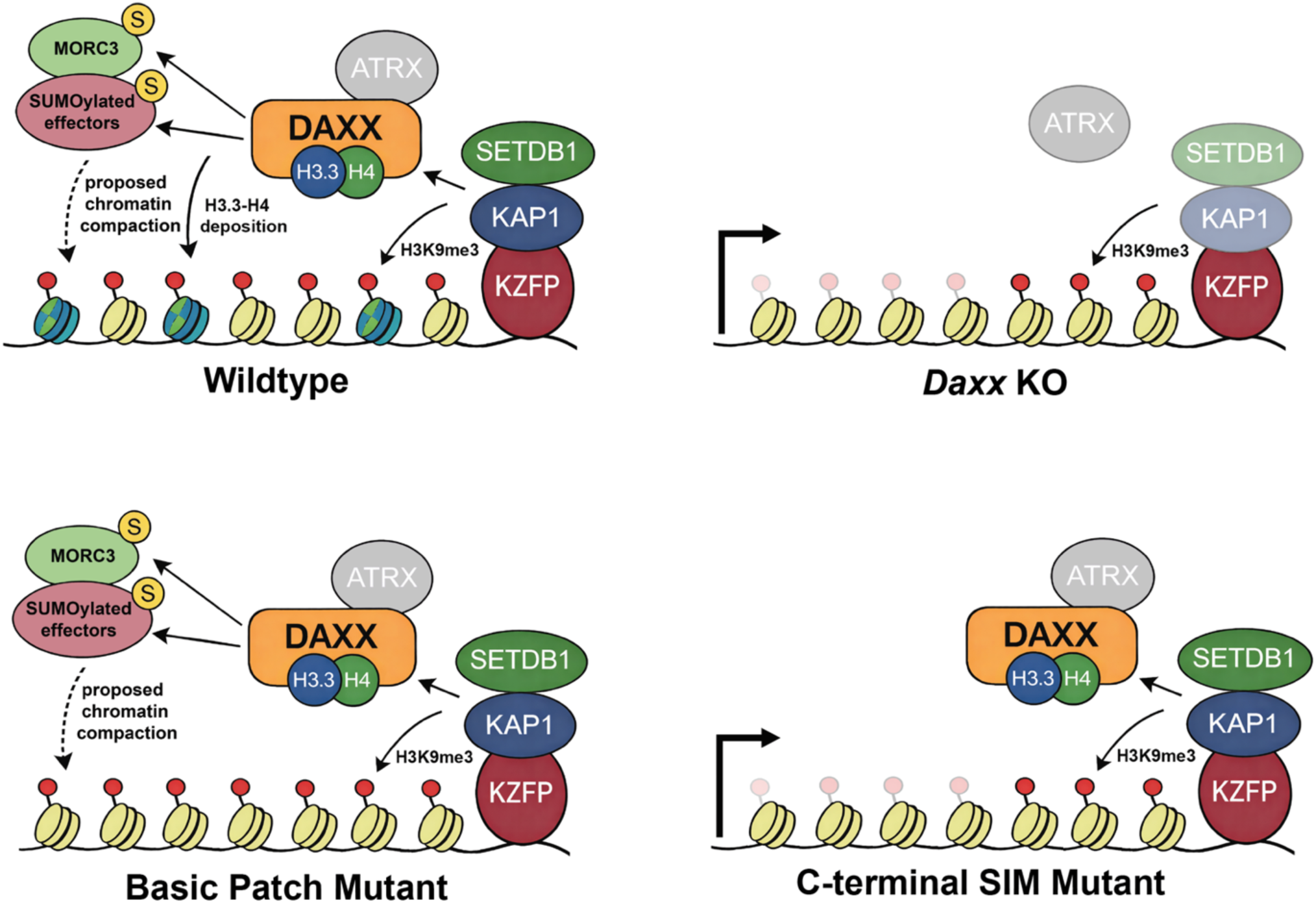
DAXX uses separable determinants for H3.3 deposition and ERV silencing, with the C-terminal SIM required for SUMO-dependent repression. Model: Following recruitment to ERVs, the DAXX basic patch promotes DNA engagement and H3.3-H4 deposition, resulting in local H3.3 enrichment. The DAXX C-terminal SIM recruits SUMOylated effectors downstream of chromatin binding to support ERV silencing and contribute to H3.3 enrichment. Wild-type DAXX supports both outputs, whereas basic patch mutants retain DAXX occupancy and silencing but fail to restore H3.3 enrichment. By contrast, C-terminal SIM mutants retain DAXX occupancy but fail to recruit MORC3 and other SUMOylated effectors, lose silencing, and show reduced H3.3 enrichment at DAXX-bound ERVs. The dashed arrow denotes a proposed MORC3-associated chromatin compaction step. ATRX is shown in gray to indicate that the ATRX-DAXX interaction is dispensable at these loci.

### A basic residue-enriched surface promotes productive H3.3 deposition through DNA engagement

The modular framework above clarifies that DAXX-directed H3.3 enrichment reflects a distinct deposition activity that can be mechanistically interrogated independently of repression output. Structures of the DAXX histone-binding domain show an H3.3-H4 heterodimer tightly wrapped and shielded from DNA, consistent with a pre-deposition complex^60–62^. A central mechanistic question has therefore been how DAXX transitions from histone shielding to a DNA-engaged intermediate that enables nucleosome assembly. Our data identify a basic patch within the tower region of the histone-binding domain that functions as a DNA engagement module, promoting DNA binding and productive nucleosome assembly *in vitro* and supporting H3.3 enrichment at endogenous DAXX targets in cells. This finding is notable in the context of prevailing models in which histone chaperones minimize nonspecific histone-DNA binding while escorting histones, relying on sequestration of histone DNA-binding surfaces to prevent aberrant histone-DNA interactions prior to histone deposition^23,32,76,77^. The data instead support a two-surface logic in which histone shielding and DNA engagement are integrated within the same chaperone, enabling a controlled transition into deposition competence.

This architecture has become increasingly recognized across histone chaperone systems. In CAF-1, a lysine-, glutamic acid, arginine-rich module and a winged helix DNA-binding domain that cooperatively support DNA engagement and deposition outcomes^64–67,69,78,79^. A related principle is evidence in a distinct yeast H3-H4 chaperone Rtt106, which contains a positively charged DNA-binding ridge, and selective disruption of this surface impairs silencing and transcriptional fidelity^80–84^. Likewise, in the HIRA complex, the UBN1 subunit uses conserved lysine residues to engage DNA directly, demonstrating that basic residues within chaperone modules can mediate DNA binding during replication-independent deposition^85–88^. Together, these examples support a general model in which localized basic surfaces on histone chaperones act as DNA capture elements that stabilize an initial chaperone-histone-DNA intermediate. Within this framework, the DAXX basic patch likely lowers the kinetic barrier for productive DNA engagement and promotes a histones handoff into a wrapped nucleosomal state.

### Recruitment versus engagement separates DAXX localization from productive H3.3 deposition at ERVs

Building on the identification of the DAXX basic patch as a DNA-binding element, our results show that, for DAXX at ERVs, chromatin recruitment can be separated from the subsequent DNA-engagement step required for productive H3.3 deposition. In pluripotent cells, these ERVs are embedded within a multicomponent KRAB zinc finger protein-TRIM28/KAP1 repressive architecture that establishes H3K9me3-marked heterochromatin at defined retrotransposon elements^56,57,89–91^. This chromatin environment provides a setting in which DAXX occupancy can be maintained even when the basic patch-dependent deposition step is compromised.

The basic patch mutant phenotypes provide direct evidence for this DAXX-specific separation of localization from deposition. DAXX occupancy at ERV-associated targets persists when the basic patch is disrupted, yet H3.3 enrichment at the same loci is reduced, indicating that stable chromatin association can be uncoupled from deposition competence. These observations refine prior work showing that DAXX contributes to ERV repression through an ATRX-independent axis that interfaces with canonical H3K9 silencing machinery^37^. More broadly, the data support a two-step model in which ERV-directed pathways establish the chromatin context for DAXX localization, whereas the basic patch mediates a post-recruitment DNA-engagement step that couples the DAXX-H3.3-H4 complex to substrate DNA and enables nucleosome assembly.

Several studies have implicated ATRX in retrotransposon silencing, motivating models in which ATRX-DAXX-mediated H3.3 deposition helps establish or maintain heterochromatin at ERVs^35,36,38,46,92,93^. These models further suggest that ATRX recruitment is promoted by H3K9me3-rich chromatin, consistent with H3K9me3 recognition by the ATRX ADD domain^94,95^. In our study, perturbations that disrupt ATRX binding, including substitutions in the DAXX four-helix bundle that comprises the ATRX-interaction surface^37,96^ did not measurably affect DAXX occupancy, H3.3 enrichment, or ERV repression at DAXX-bound ERVK loci. These results align with prior structure-guided disruption of the same interface, which uncoupled ATRX-DAXX complex formation from DAXX-dependent retrotransposon silencing^37^. Collectively, these findings support a repeat-class and locus-specific requirement for ATRX-DAXX, consistent with evidence that the complex becomes more stringently required at other repeat compartments, including tandem repeats under conditions of DNA hypomethylation^36^.

### H3.3 deposition at DAXX-bound ERVs is dispensable for silencing in mESCs

Defining a post-recruitment DNA-engagement step provides a framework to ask whether DAXX-directed H3.3 deposition is itself required for repression or instead represents a separable chromatin output at DAXX-bound ERVs. Although H3.3 enrichment at ERVs has been associated with ERV control and silent chromatin features in embryonic stem cells, these correlations do not establish that stable incorporation of H3.3-containing nucleosomes is required for silencing. In our previous work^37^, we addressed this question directly at DAXX-bound ERVs in mESCs. We found that H3.3-knockout mESCs exhibit markedly reduced DAXX steady-state abundance, and re-expression of wild-type H3.3 restores DAXX levels. We further found that an H3.3(L126A/I130A) allele, which retains binding to DAXX but disrupts the tetramerization interface required for stable chromatin incorporation, stabilizes DAXX yet is not detectably incorporated at ERV LTRs. Despite the absence of detectable H3.3 deposition, silencing of DAXX-regulated ERV families was maintained. These findings support a model in which soluble H3.3-H4 sustains DAXX abundance and silencing competence, whereas stable H3.3 deposition at these loci is dispensable, consistent with evidence that histone engagement stabilizes DAXX histone-binding domain^43,62^.

Our study extends this separation to DAXX itself by uncoupling chromatin recruitment from productive deposition. DAXX basic patch mutants remain recruited to ERVs and retain histone association, yet they fail to restore H3.3 enrichment at DAXX-bound elements, consistent with a post-recruitment defect in substrate DNA engagement and histone transfer. Despite this deposition defect, DAXX-mediated ERV silencing remains intact. Together, the incorporation-defective H3.3(L126A/I130A) allele and the DAXX basic patch separation-of-function mutants indicate that H3.3 enrichment is dispensable for repression of DAXX-regulated ERVs and ERV-linked transcripts in mESCs. We therefore favor a model in which DAXX-directed H3.3 deposition shapes chromatin properties, including nucleosome organization, turnover, and longer-term epigenetic stability, rather than serving as the primary determinant of transcriptional repression.

### DAXX C-terminal SIM couples effector engagement to ERV silencing and to H3.3 enrichment

Another clear separation between recruitment and downstream function is revealed by disruption of the DAXX C-terminal SIM. The DAXX CSM protein retains occupancy at ERV-associated loci, yet fails to restore local H3.3 enrichment and fails to silence ERV transcription. This genetic ordering places the SIM downstream of chromatin recruitment and upstream of an effector engagement step that supports both ERV silencing and H3.3 enrichment, consistent with a role for SIM-mediated interactions with SUMOylated partners. The DAXX C-terminal SIM mediates association with SUMO-modified PML and localization to PML nuclear bodies, establishing precedent for SIM-dependent engagement of SUMOylated scaffolds and compartmentalized silencing ^97–100^.

These observations suggest that the DAXX C-terminal SIM functions as a post-recruitment effector interface that can engage SUMOylated factors within ERV-associated heterochromatin, rather than serving exclusively as a MORC3 docking site. TRIM28/KAP1 undergoes extensive SUMOylation, and SUMO-dependent interactions promote recruitment of SETDB1 together with chromatin remodeling and deacetylase activities, providing plausible routes through which SIM-mediated contacts could stabilize local repressor architecture at ERVs^101–103^. Within this network, MORC3 is a SUMOylated MORC-family ATPase that binds DAXX and contributes to silencing of selected ERV families in mESCs^46^. We find that DAXX CSM mutants lose association with MORC3 and fail to recruit MORC3 to ERV loci, while retaining interactions with TRIM28/KAP1 and SETDB1. These results support a selective defect in downstream effector engagement rather than a generalized loss of repressor complex assembly. Conversely, MORC3 depletion reduces H3.3 enrichment at DAXX-bound peaks, linking SIM-dependent effector engagement to the H3.3 enrichment phenotype at ERV loci. Together, these observations support a model in which the DAXX C-terminal SIM enables a SIM-dependent effector step that includes MORC3 and may also involve additional SUMOylated scaffold or effector proteins.

### MORC-family DNA compaction provides a plausible effector mechanism for MORC3-dependent silencing

The genetic separations described above raise the question of which SIM-engaged effector activities enforce ERV silencing, and MORC-family DNA compaction offers one plausible answer. Biochemical analysis of MORC2 revealed ATP hydrolysis-dependent DNA compaction *in vitro*, whereas *C. elegans* MORC-1 topologically entraps DNA and compacts it through loop-forming multimeric assemblies^104,105^. These mechanistic precedents support a working model in which MORC3 acts downstream of DAXX recruitment through ATPase-driven DNA engagement and loop trapping that locally constrains chromatin over ERV regulatory sequences. Such compaction would be expected to disfavor productive transcription and to oppose chromatin-remodeling activities that promote nucleosome eviction. At the same time, by stabilizing a compacted substrate and limiting DNA mobility, MORC3 could increase the likelihood that transiently exposed DNA is captured by DAXX-H3.3-H4, thereby elevating H3.3 occupancy as a coupled consequence of effector action rather than as the primary determinant of silencing. This model is consistent with evidence that MORC3 contributes to repeat silencing pathways associated with reduced accessibility and heterochromatin features^10^.

### A modular DAXX pathway for repeat control and genome stability

The modular architecture defined here positions DAXX as an interaction hub that couples ERV recruitment platforms to separable downstream outputs. One output is basic patch-mediated DNA engagement that enables productive H3.3 nucleosome assembly. A second output is a SUMO-dependent effector engagement within a SUMO-rich repression environment, potentially involving PML, TRIM28/KAP1-associated complexes, and MORC3. This framework provides a basis for understanding how distinct DAXX perturbations may differentially impact histone deposition, repeat control, and genome stability across cellular contexts.

These considerations are likely to extend to disease settings in which the DAXX pathway is recurrently altered. Pancreatic neuroendocrine tumors frequently harbor *DAXX* and *ATRX* alterations and are associated with alternative lengthening of telomeres (ALT), linking disruption of repeat-associated chromatin regulation to genome instability phenotypes^48,49^. ATRX-DAXX pathway perturbation and/or ALT have been reported in additional tumor types, including neuroblastoma, IDH-mutant diffuse gliomas, osteosarcoma, and uterine leiomyosarcoma^106–110^. More recent functional analyses indicate that cancer-associated variants affecting the DAXX C-terminal SUMO-interacting motif can compromise repeat-associated outputs, including transcriptional repression^111^. Together, these observations motivate future studies that map how discrete DAXX modules, including the basic patch and C-terminal SUMO-interacting motif, contribute to repeat regulation in tumor lineages and to the broader epigenetic consequences of DAXX dysfunction.

In summary, DAXX recruitment to ERVs can be uncoupled from two downstream activities: DNA engagement that supports H3.3 deposition, and SUMO-dependent effector engagement that enforces silencing. The separation-of-function mutations defined here enable mechanistic dissection of how histone chaperone activity and SIM-dependent pathways are coordinated to maintain repeat silencing and genome stability. By partitioning recruitment, DNA engagement, and effector coupling into discrete modules, this work provides a testable framework for linking ATRX-DAXX pathway lesions to repeat dysregulation and genome instability in development and cancer.

## METHODS

### ES cell lines and culture

The *Daxx* ^−/−^ mESCs were provided by Philip Leder^112^. mESCs were maintained under 2i + LIF conditions in a medium containing a 1:1 mix of Dulbecco’s modified Eagle’s medium (DMEM)/F12 (Life Technologies 11330-033) and ATCC mouse ES cell basal medium (Fisher Scientific 50-238-3794) with the addition of N2 and B27 supplements (Life Technologies 17502-048 and 17502-044, 1:100 dilutions), penicillin/streptomycin (100 U/ml and 100 µg/ml, respectively; Life Technologies 15140-122), 0.1 mM 2-mercaptoethanol, 1X GlutaMAX^TM^-I (Life Technologies 35050-061), 1X MEM NEAA (non-essential amino acids; Life Technologies 11140-050), Leukemia inhibitory factor (LIF), CHIR99021 at 3 μM (TOCRIS) and PD0325901 at 1 μM (Selleckchem). mESCs were grown in feeder-free conditions on dishes coated with 0.1% bovine gelatin.

### Immortalized cell lines and culture

MEFs (*Daxx* ^−/−^) were provided by Philip Leder^112^. MEFs and 293T (ATCC CRL-3216) were maintained in DMEM supplemented with 1X GlutaMAX^TM^, penicillin/streptomycin, and 10% fetal bovine serum (Gibco).

### Lentiviral plasmid construction, lentiviral production, and transduction

HA-tagged H3.3 and FLAG-tagged murine DAXX (mDAXX) or human DAXX (hDAXX) were cloned into pCDH-EF1-MCS-Puro lentiviral vectors (System Biosciences). 293T cells were transfected with lentiviral vector and helper plasmids (psPAX2 and pMD2.G) to produce lentiviruses. The supernatant containing lentiviruses was collected, filtered, and concentrated after 72 h. To generate cell lines stably expressing various mDAXX transgenes, cells were transduced with concentrated lentiviruses in the presence of 5 µg/ml Polybrene (Santa Cruz Biotechnology sc-134220). Transduced cells were grown under puromycin selection (1 µg/ml for mDAXX transgenes) 48 h after transduction. Cells were collected for downstream experiments within 1–3 weeks after lentiviral transduction.

### Cycloheximide chase assay

MEF *DAXX −/−* rescued with wildtype, BPM, or HBM DAXX transgenes were treated with 10 µg/ml of cycloheximide. The treated cells were collected at 0, 10, and 24h post-treatment to analyze the turnover rate of DAXX proteins.

### Reverse transcription and quantitative PCR

RNA was extracted using a Quick-RNA MiniPrep kit (Zymo), digested with DNase I (Turbo) off-column, followed by a second column clean-up. cDNA was prepared from 1 µg RNA using the ProtoScript II First Strand cDNA Synthesis (New England Biolabs; random hexamer primers). Quantitative real-time PCR was performed on diluted cDNA in the presence of 0.5 µM forward and reverse primers using Power SYBR Green Master Mix. Program: Initial 95 °C for 10s, then 95 °C for 20s, 60 °C for 30s, 72 °C for 20s (52 cycles).

### Immunoblots

Whole-cell lysates or immunoprecipitation samples were run on SDS-PAGE gels, transferred to nitrocellulose membrane, blocked in 5% non-fat milk in Tris-buffered saline with 0.5% Tween-20, probed with primary antibodies, and detected with horseradish peroxidase-conjugated anti-rabbit or anti-mouse secondary antibodies (Bio-Rad).

### Chromatin immunoprecipitation

Approximately 30 million mESCs were crosslinked with 1% Paraformaldehyde (Electron Microscopy Sciences) for 10 min at 37 °C and quenched with 200 mM glycine. To isolate nuclei, cells were resuspended in lysis buffer (50 mM HEPES pH 7.9, 140 mM NaCl, 1 mM EDTA, 10% glycerol, 0.5% NP40, 0.25% Triton X-100). Nuclei were washed with digestion buffer (50 mM HEPES pH 7.9, 1 mM CaCl_2_, 20 mM NaCl, 1x protease inhibitor cocktail, and 0.5 mM PMSF), and treated with 80 Units of MNase (Worthington Biochemical Corporation, LS004798) for 10 min at 37 °C. Digestion reaction was quenched by adding 10 mM EDTA, 5 mM EGTA, 80 mM NaCl, 0.1% sodium deoxycholate, and 0.2% SDS. Nucleosomes were further solubilized by sonication using Covaris S220 (160 peak incidental power, 5% duty factor, 200 cycles/burst, 45” on −30” off) 7-times. Chromatin was supplemented with 1% Triton X-100, and insoluble chromatin was removed by centrifugation. Chromatin concentration was measured using qubit, and spike-in chromatin (chromatin from K562 cells overexpressing FLAG-tagged hDAXX and HA-tagged H3.3 prepared in the same way) was added at a 1:50 ratio. 25 µg chromatin was incubated with primary antibodies overnight for ChIP: DAXX (FLAG 1:50), H3.3 (HA 1:75), KAP1 (1:50), and MORC3 (1:50). Primary antibodies were captured using Dynabeads and washed 3 times with RIPA buffer, 2 times with RIPA + 300 mM NaCl, and 2 times with LiCl buffer. Chromatin was reverse-crosslinked at 65 °C overnight in buffer containing 10 mM Tris, 1 mM EDTA, and 1% SDS. Chromatin was then incubated with RNase A for 1h at 37 °C and proteinase K for 2h at 55 °C and DNA was subsequently purified using PCR purification columns. Eluted DNA was diluted 1:15 for qPCR analysis. Paired-end HT-Sequencing libraries were prepared using NEBNext Ultra II DNA Library Prep Kit for Illumina with NEBNext Multiplex Unique Dual Oligos. Sequencing was performed on a NovaSeq6000 at the UW Biocore Sequencing Facility. ChIPs were performed in at least two biological replicates.

### DAXX HBD-H3-H4 complex /H3-H4 tetramers purification

Human DAXX fragment (aa 183–417) was cloned into pET28b to generate an N-terminal His_6_-FLAG-DAXX HBD (histone binding domain). Human histone H3.3 and codon-optimized H4 were cloned into pRUTH5 to yield an N-terminal His_6_-TEV tag (courtesy of A. Ruthenburg). Single or multiple point mutations were introduced by standard mutagenesis PCR procedures. Histones and DAXX HBD were individually expressed in BL21 Star (DE3) cells (Invitrogen) into inclusion bodies for 4 h at 37 °C. Inclusion bodies were resolubilized in 6 M guanidine-HCl, 1 M NaCl, 25 mM Tris-HCl pH 8, and purified on a Ni-NTA affinity column. To prepare H3–H4 tetramers, equimolar ratios of H3 and H4 were mixed in refolding buffer (4 M guanidine, 25 mM HEPES pH 7.9, 1 M NaCl, 5 mM EDTA, 15% glycerol) and dialyzed against 25 mM HEPES pH 7.9, 1 M NaCl, 1 mM EDTA, 10% glycerol. To prepare HBD-H3–H4 complexes, equimolar ratios of HBD, H3, and H4 were mixed in refolding buffer (4 M guanidine, 25 mM HEPES pH 7.9, 1 M NaCl, 5 mM EDTA, 15% glycerol) and dialyzed against 25 mM HEPES pH 7.9, 1 M NaCl, 1 mM EDTA, 10% glycerol. The resulting protein complexes were further purified on a HiLoad 16/600 Superdex 200 column (GE Healthcare).

### DAXX (full length)-H3-H4 complex purification

Murine *DAXX* (wildtype or mutant) was cloned into pCAG vector to yield an N-terminal FLAG-DAXX. Human H3.3 was cloned into pCAG to yield a C-terminally strepII-tagged H3.3. Human H4 was cloned into pCAG. The three plasmids were co-transfected in equimolar amounts into 293T *DAXX −/−*. 60 h post-transfection, cells were homogenized in hypotonic lysis buffer (20 mM HEPES pH 7.9, 10mM KCl, 5mM MgCl2, 0.5mM EGTA, 0.1mM EDTA 1mM DTT, 1mM Benzamidine, 0.8mM PMSF) and lysed in the buffer (20mM HEPES pH 7.9, 110mM KCl, 2mM MgCl_2_, 0.1mM EDTA, 1mM DTT, 1x Protease inhibitor cocktail, 0.4mM PMSF). The resulting lysate was incubated in 400 mM ammonium sulfate and spun down in ultracentrifuge (28.5Kxg for 1.5h). Nuclear extract was dialyzed against buffer (20mM HEPES pH 7.9, 250mM KCl, 1mM EDTA, 0.01% NP-40, 0.4mM PMSF, 2mM BME) for 2h twice. Nuclear extract was incubated with Strep-TactinXT 4Flow (IBA) for 2h. Beads were washed 3 times with the buffer (15 mM HEPES pH 7.9, 750 mM KCl, 1 mM EDTA, 0.05% Triton X-100, 8 mM PMSF) and captured proteins were eluted with 50 mM D-(+)-Biotin (Enzo). The purified complex was subsequently incubated with M2 anti-FLAG affinity gel (Sigma A2220) for 3h. Beads were washed 3 times with the buffer (15 mM HEPES pH 7.9, 750 mM KCl, 1 mM EDTA, 0.05% Triton X-100, 8 mM PMSF), and captured proteins were eluted using 300 µg/ml of 3xFLAG peptide. The protein complex was further purified on Mono Q PC 1.6/5 column (GE Healthcare). Elution fractions were analyzed using SDS-PAGE, followed by staining with Coomassie Brilliant Blue.

### FLAG co-immunoprecipitation

Cell lysate was prepared by resuspending approximately 40 million MEFs (with stable transgenes) in a lysis buffer (20 mM HEPES pH 7.9, 300 mM KCl, 1 mM EDTA, 0.12% Triton X-100, 2 mM 2-mercaptoethanol, 0.4 mM PMSF, 2x protease inhibitor cocktail), followed by douncing and separation of the insoluble fraction by centrifugation. Per sample, 40–60 µl of packed anti-FLAG M2 beads (Sigma) were added to the lysate and incubated at 4 °C for 3h. Beads were washed three times with wash buffer (same as lysis buffer without 2-mercaptoethanol) for 5 min each. Finally, beads were transferred into microspin columns and samples were eluted with 30–60 µl elution buffer (wash buffer supplemented with 0.5 µg/µl 3×FLAG peptide) after 5 min incubation on ice.

### Electrophoretic mobility shift assay

Cy5-labelled widom601 DNA was used to assay protein-DNA binding. Protein complex-DNA binding reactions were performed in the buffer (50 mM Tris pH 7.6, 200 mM KCl, 0.025% NP-40, 2 mM MgCl_2_, 0.1 mg/mL BSA, 10% glycerol, 0.4 mM PMSF, and 1 mM DTT). Incubations were done for 45 minutes at 30°C and ran on a 0.7% native agarose gel prepared in 1X TBE at 6 V/cm for 60 minutes in a cold room. Fluorescent signals were imaged with Typhoon 5. Signal intensities for bound and unbound fractions were quantified by measuring integrated density signals with ImageJ. % bound entities from three independent experiments were plotted in GraphPad Prism 10 software, and apparent K_D_ values were obtained by fitting the data to a four-parameter Hill curve with variable slope (specific binding) with the equation:

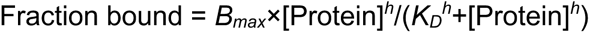

where *B*_max_ corresponds to maximum binding, *h* represents the Hill coefficient, and *K*_D_ refers to the dissociation constant.

### Plasmid DNA supercoiling assay

Plasmid pCRBlunt (Invitrogen) was relaxed with topoisomerase I at 30 °C for 30 min in the buffer (10 mM Tris pH 7.6, 40 mM KCl, 2 mM MgCl_2_, 60 mM NaCl, 0.5 mM EGTA, 1 mM DTT, 0.2 µg/ml BSA), 250 ng per reaction. Pre-relaxed DNA was incubated with 0.6 μM or 1.2 μM of histones or chaperone-histone complexes at 37 °C for 1h in a buffer (10 mM Tris pH 7.6, 50 mM NaCl, 2 mM MgCl_2_, 0.5 mM DTT, 25 ng/ml BSA, 0.5 mM EDTA, 0.01% NP40, 0.2% glycerol). The reaction was quenched by adding 0.3% SDS and 0.5 µg Proteinase K and incubating at 55 °C for 20 min. The product was cleaned up with phenol chloroform extraction. Approximately 150 ng of reaction product was run on a 1.2 % agarose gel in 1× TAE buffer at 50V for 14h and stained with ethidium bromide post-run.

### Bioinformatics and software used in the study

#### ChIP-seq

Reads were aligned to mouse (mm10) or human (hg38) genomes using bowtie2 (parameters: -q -v 2 -m 2 -a – best –strata)^113^. Sample normalization factor was determined as ChIP-Rx = 10^^6^ / (total reads aligned to exogenous reference genome) or RPKM = 10^^6^ / (total aligned reads). Samtools was used to convert sam files to bam files^114^. Deeptools was used to generate bigwig files^115^. Peaks were called using MACS2^116^. Peak annotation was done with HOMER^117^. Deeptools^118^ and IGV genome browser were used for data visualization. The age of transposable elements was estimated based on sequence divergence, assuming a neutral mutation accumulation rate of 0.2% per million years^119,120^.

#### RNA-seq

Reads were aligned to mouse (mm10) transcriptome using STAR −2.7.8a using the following parameters --runThreadN 32 --quantMode TranscriptomeSAM GeneCounts --outSAMtype BAM SortedByCoordinate)^121^. Read counts for genes and repeats were derived using TEtranscripts and normalized using DESeq2^122^. Differentially expressed genes and repeats were determined using DESeq2 (adjusted *p*-value < 0.05)^123^. ggplot2 was used to visualize data^124^. The distance of differentially expressed genes from DAXX peaks was estimated using bedtools^125^.

#### Others

AlphaFold2 was used to predict 3D interaction of DAXX HBD with widom601 DNA and to draw a comparison of MORC3 and MORC2 ATPase domains^126^.The AlphaMissense database was used to predict missense variant effect on all the residues of human DAXX protein^70^. The working model was created with BioRender.com. The sequence alignment comparing the DAXX amino acid sequence across different vertebrate species was performed with Snapgene, using the Clustal Omega algorithm. Tumor sample genome sequencing was visualized using SnapGene^71^.PhosphoSitePlus was used to predict potential phosphorylation sites in the mouse DAXX protein^127^.

#### Publicly available datasets used in this study

For DAXX HBD-H3.3-H4 3D structure (**Fig. 2A**), PDB 4H9N was used. For MORC3 ChIP-MS (**Fig. S5E**), PXD027368 was used. For MORC3 ChIP-seq (**Fig. S1E, S5G**), H3.3 ChIP-seq in MORC3 KO background (**Fig. S5D, 4H-I**), MORC3 KO and WT RNA-seq data (**Fig. S7F**), GSE159936 was used. For DAXX KO and WT mESCs RNA-seq (**Fig. 1D-F, S7F**), GSE102688 was used. For PML RNA-seq (**Fig. S7F**), E-MTAB-10153 was used. For ATRX ChIP-seq (**Fig. S1E**), GSE151054 was used. For SMARCAD1 ChIP-seq (**Fig. S1E**), E-MTAB-7012 was used. For KAP1 ChIP-seq (**Fig. S1E**), GSM1555120 was used. For H4K20me3 ChIP-seq (**Fig. S1E**), GSM307622 was used. For H3K27me3 ChIP-seq (**Fig. S1E**), GSM1033638 was used. For SETDB1 ChIP-seq (**Fig. S1E**), GSM459273 was used. For H3K9me3 ChIP-seq (**Fig. S1E**), E-MTAB-7012 was used.

## Data availability

ChIP-sequencing data generated in this study are available on the NCBI Gene Expression Omnibus database with accession number:

## Acknowledgements

We thank Melissa Harrison and Siddhant Jain for comments on the manuscript. P.W.L. acknowledges support from the Neuroendocrine Tumor Research Foundation and the Pew Scholars Program in the Biomedical Sciences. D.H. was supported by a Boehringer Ingelheim Fonds PhD fellowship. This work was supported by NIH grants P01CA196539 and R01CA266861.

---

**Figure S1.**
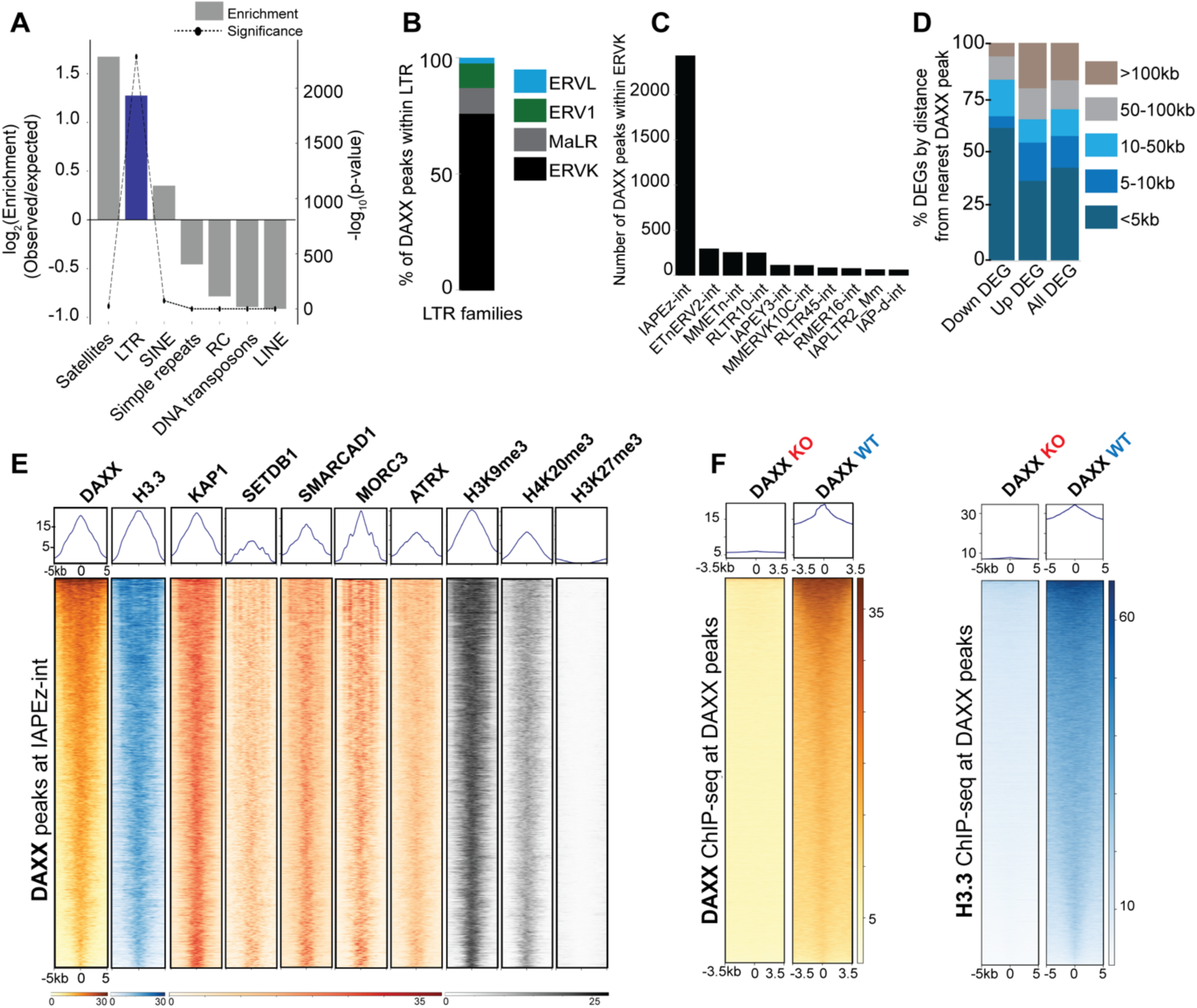
DAXX occupancy is enriched at ERVK elements and correlates with local H3.3 enrichment in mESCs. (**A**) Enrichment of DAXX peaks across major repeat classes (log_2_ observed/expected; dashed line, −log_10_ P value from overlap test). RC, rolling-circle elements. (**B**) Composition of LTR-overlapping DAXX peaks by LTR class (ERVK, MaLR, ERV1, ERVL). (**C**) DAXX peak counts across ERVK subfamilies, highlighting predominant overlap with IAPEz-int and a limited set of additional lineages, including ETnERV2-int. (**D**) Differentially expressed genes (DEGs) in *Daxx* knockout versus wild-type rescue mESCs binned by distance to the nearest DAXX peak; “Up” and “Down” indicate higher or lower expression in knockout. (**E**) Metaprofiles and heat maps of the indicated ChIP-seq signals centered on DAXX peak summits within IAPEz-int elements (±5 kb), with peaks ordered identically across tracks. (**F**) Metaprofiles and heat maps of H3.3 (±5 kb) and DAXX (±3.5 kb) ChIP-seq at DAXX peaks in *Daxx* knockout and wild-type rescue mESCs, showing reduced H3.3 enrichment upon DAXX loss with restoration upon rescue.

**Figure S2.**
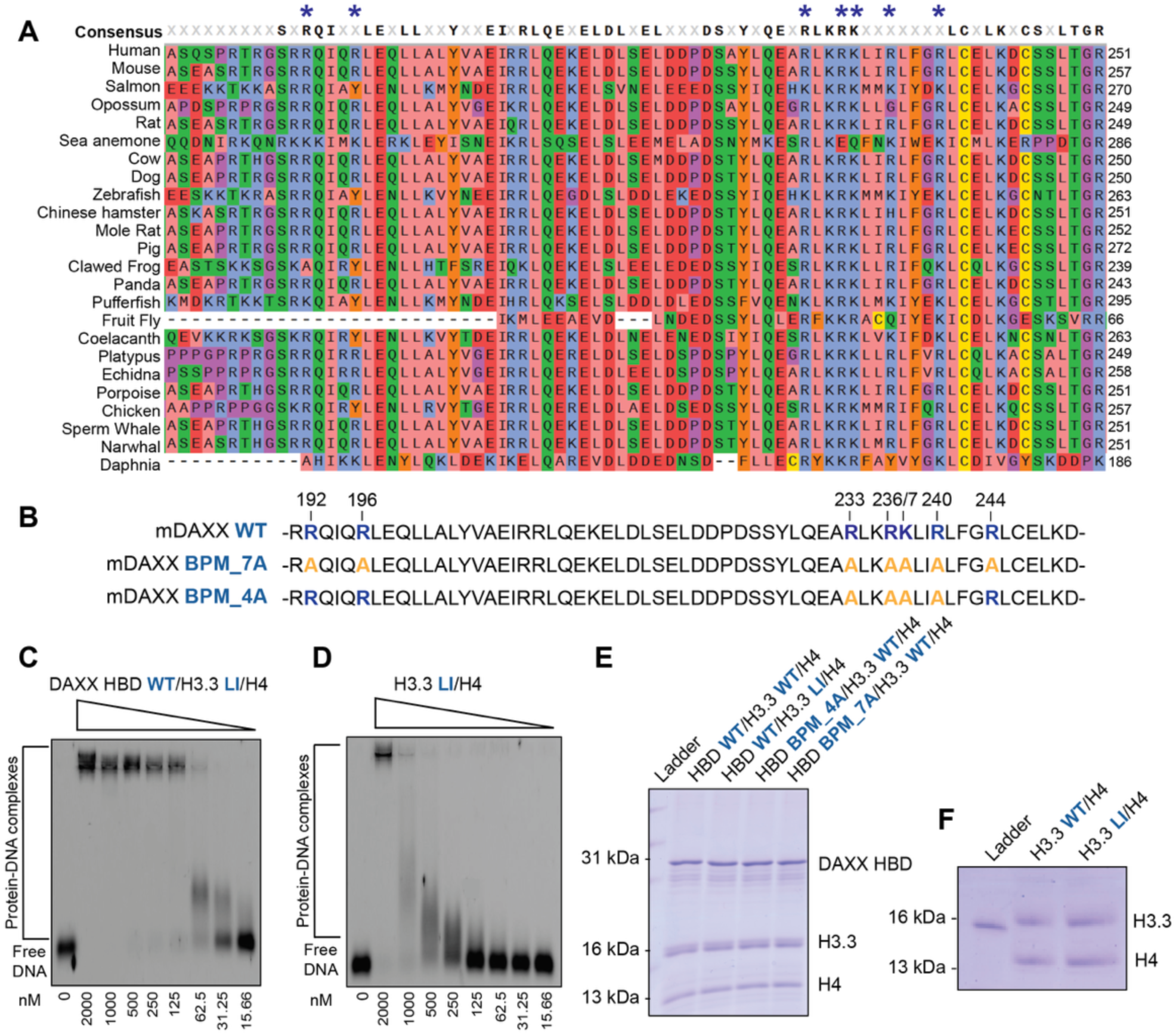
Conservation and biochemical validation of DAXX HBD basic patch mutants and EMSA reagents. **(A)** Alignment across DAXX HBD segment, showing conservation and clustering of the basic residues targeted in this study. (**B**) Basic patch substitution series in mouse DAXX HBD. (**C,D**) Representative EMSAs with Cy5-labeled DNA (approximately 7.8 nM to 2 μM titration) using H3.3(L126A,I130A)–H4 alone or DAXX HBD–H3.3(L126A,I130A)–H4, showing enhanced DNA shifting upon addition of DAXX HBD. (**E**) Coomassie-stained SDS-PAGE of purified DAXX HBD–H3.3–H4 complexes, including BPM variants, supporting preserved complex formation and stoichiometry. (**F**) Coomassie-stained SDS-PAGE of histone dimers used for EMSAs, including wild-type H3.3–H4 and H3.3(L126A,I130A)–H4, showing comparable purity.

**Figure S3.**
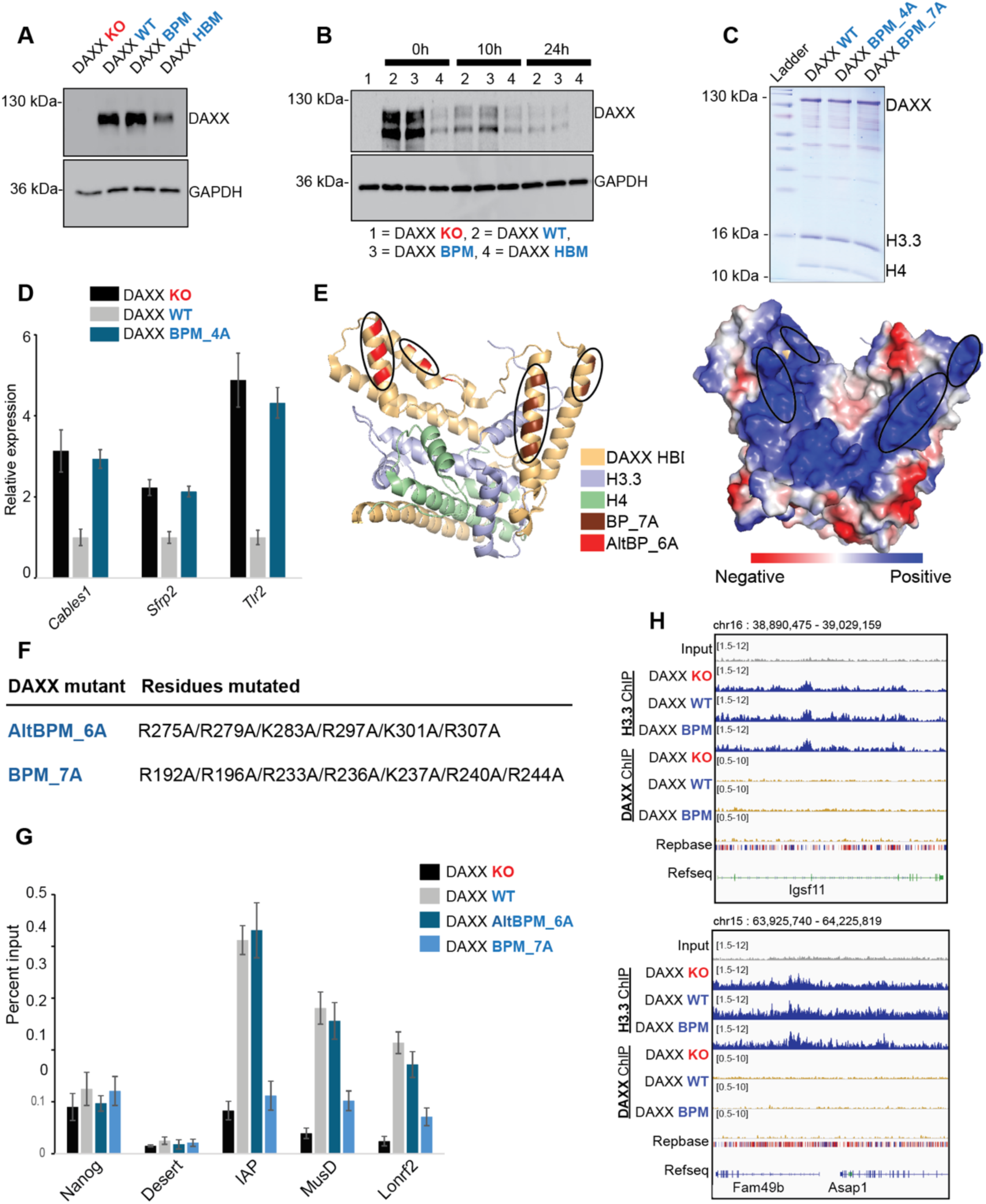
Supporting assays validate DAXX basic patch mutant phenotypes. (**A**) Immunoblot analysis of DAXX protein abundance in *Daxx-*knockout mESCs and matched rescue lines expressing wild-type DAXX, the basic patch mutant (BPM), or the histone-binding mutant (HBM). (**B**) Cycloheximide chase (0 h, 10 h, 24 h) showing similar stability of wild-type and mutant DAXX transgenes. (**C**) Coomassie-stained SDS-PAGE of purified full-length DAXX–H3.3–H4 complexes (WT, BPM_4A, BPM_7A) used in Figure 3A. (**D**) RT-qPCR of selected H3.3-dependent, DAXX-regulated genes in rescue lines, normalized to *Actb* (mean ± SEM, n = 3). (**E**) DAXX HBD electrostatic surface highlighting the tested basic patch (BP_7A) and the alternative basic surface (AltBP_6A). (**F**) AltBP_6A substitutions shown alongside BP_7A for comparison. (**G**) ChIP-qPCR for H3.3 enrichment relative to input at representative ERV loci. (**H**) Genome browser views showing H3.3 profiles at DAXX-independent loci across conditions.

**Figure S4.**
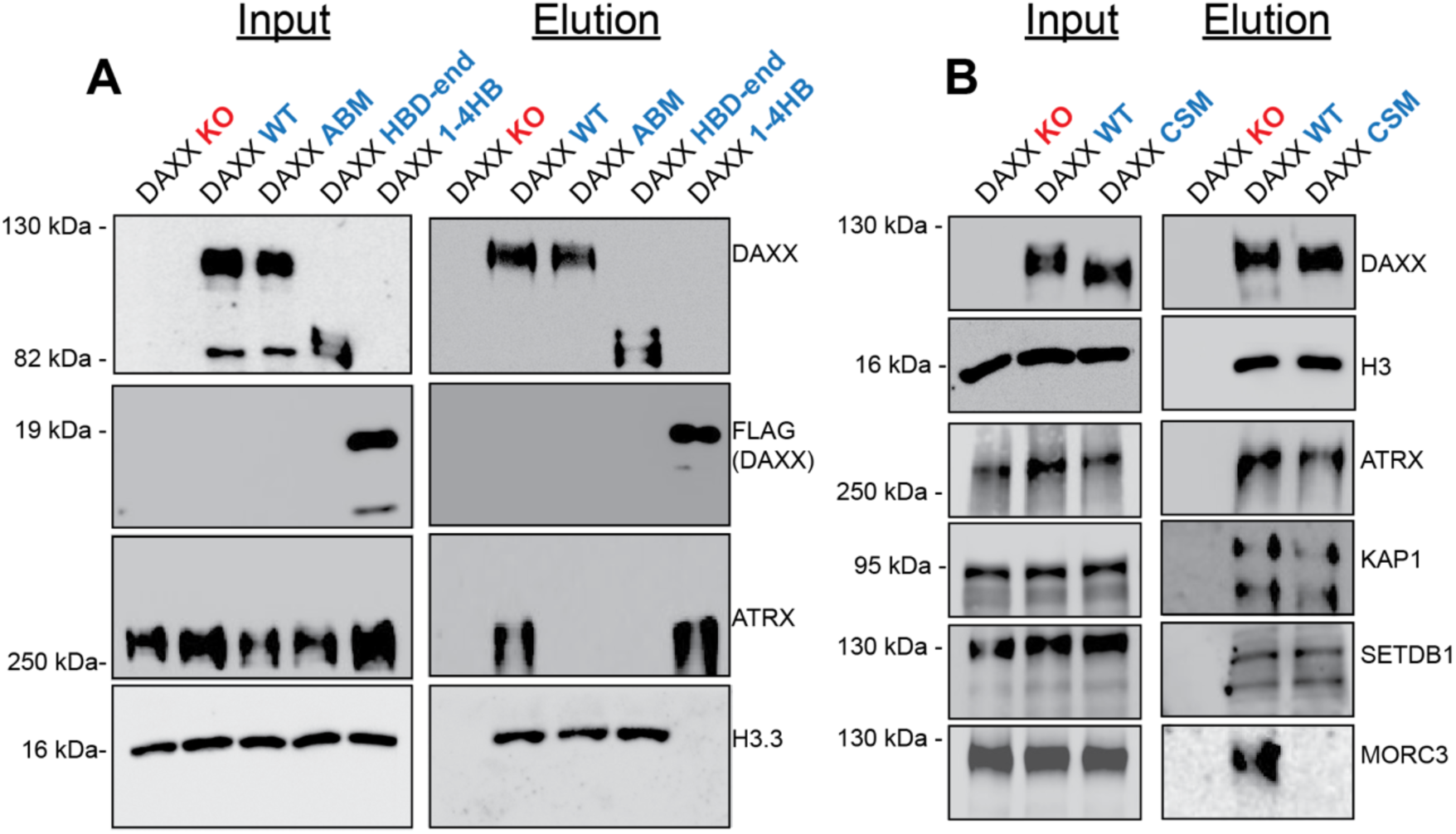
DAXX engages ATRX and MORC3 through separable interfaces. (**A**) Anti-FLAG immunoprecipitation showing ATRX co-purification with wild-type DAXX but not with Y130A (ABM), with preserved H3.3 association. Truncation controls: HBD-end enriches H3.3 with minimal ATRX, whereas 1–4HB enriches ATRX with minimal H3.3. **(B)** Anti-FLAG immunoprecipitation showing reduced MORC3 recovery with the C-terminal SIM mutant (CSM), while histone H3 and other chromatin factors (ATRX, KAP1, SETDB1) remain detectable.

**Figure S5.**
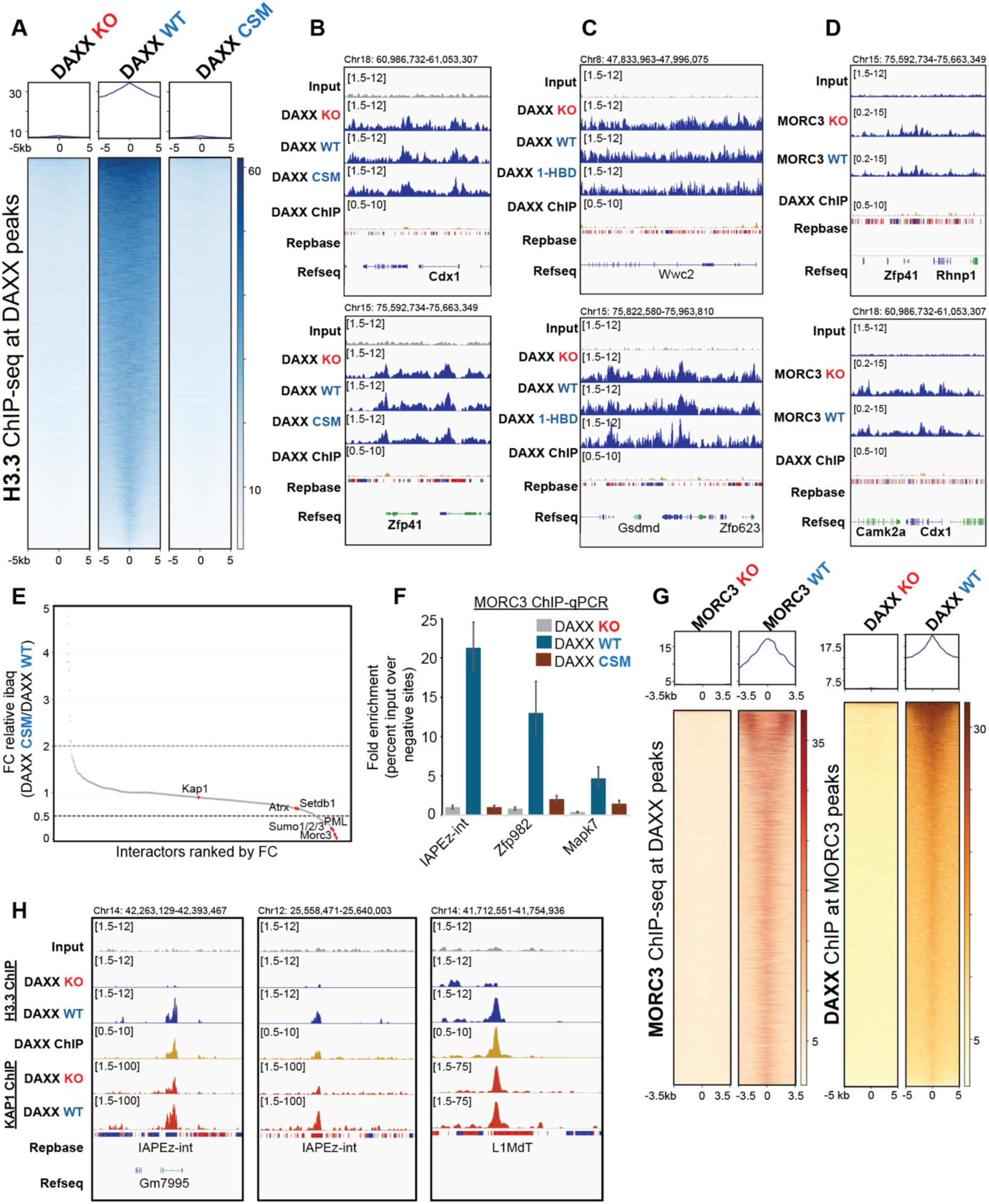
The DAXX C-terminal SIM is required for MORC3 occupancy and H3.3 enrichment at DAXX sites. **A**) H3.3 ChIP-seq at DAXX peaks, showing loss in *Daxx*-knockout cells, restoration by wild-type DAXX rescue, and failure to restore by CSM rescue. (**B–D**) Genome browser views at DAXX-independent loci showing largely unchanged H3.3 profiles across conditions. (**E**) ChIP-MS (Groh, 2021)^46^ showing reduced recovery of SUMO1/2/3 and SUMOylated interactors (including MORC3 and PML) with DAXX CSM relative to wild-type DAXX (fold change of iBAQ values). (**F**) MORC3 ChIP-qPCR at representative ERV loci in *Daxx*-knockout, wild-type rescue, and CSM rescue mESCs, showing DAXX- and SIM-dependent MORC3 occupancy. (**G**) ChIP-seq validation heat maps at DAXX peaks, showing DAXX enrichment in wild-type rescue relative to *Daxx*-knockout and MORC3 enrichment in Morc3 wild-type relative to *Morc3*-knockout. (**H**) Genome browser views at DAXX-enriched loci showing reduced KAP1 occupancy in *Daxx*-knockout cells at representative IAPEz-int targets, with an L1 locus shown for comparison.

**Figure S6.**
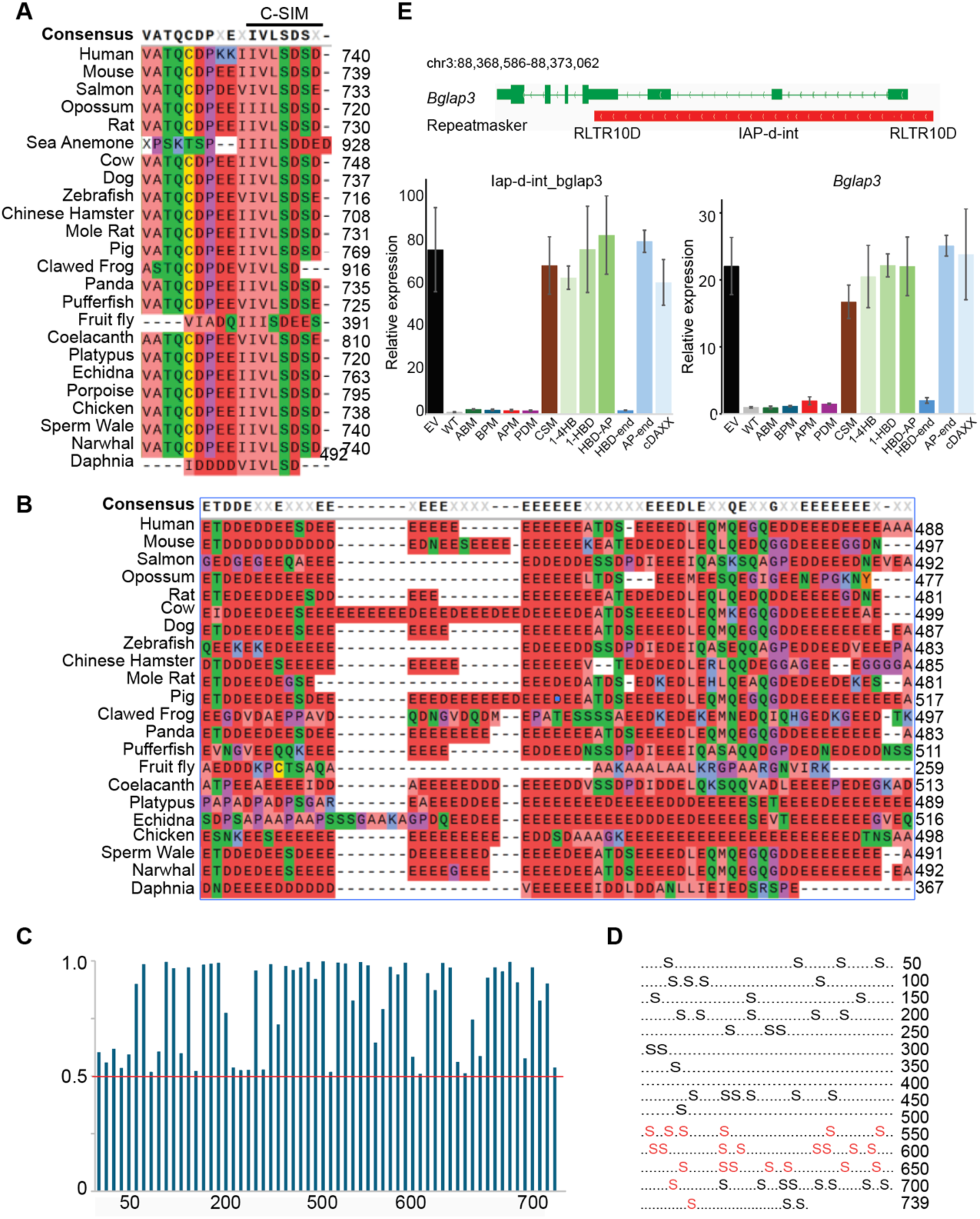
Conservation and validation of DAXX C-terminal features and domain-mapping constructs. (**A,B**) Vertebrate alignments highlighting conservation of the C-terminal SIM and biochemical conservation of the acidic patch. (**C,D**) Phosphosite prediction summaries and map of residues mutated to generate the phospho-deficient mutant. (**E**) RT-qPCR at the *bglap3* locus and its internal IAP-d-int element, showing induction in *Daxx* knockout relative to wild-type rescue, consistent with prior reports of KAP1 and SETDB1 regulation^13,45,66,67^.

**Figure S7.**
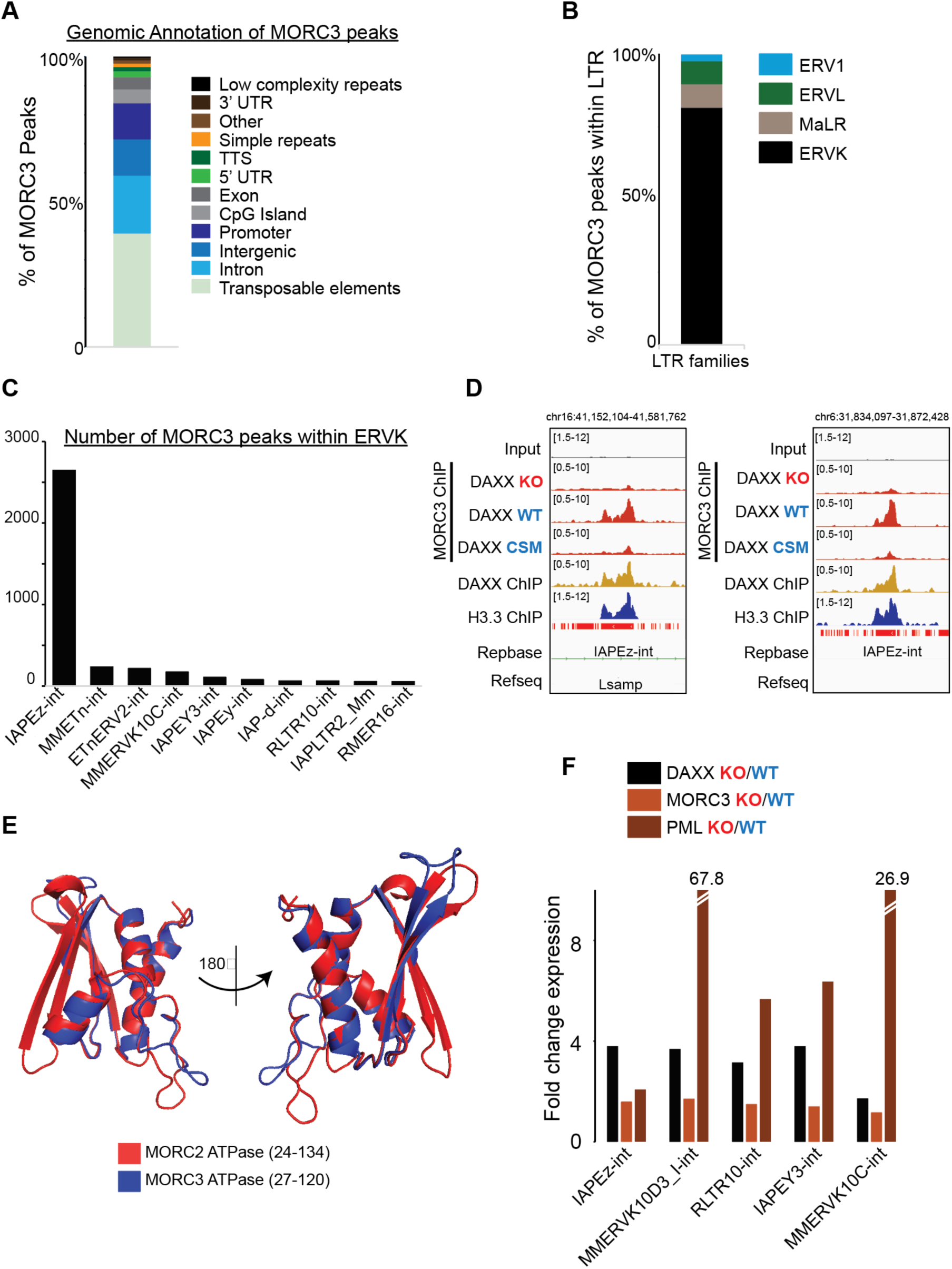
MORC3 co-targets a restricted set of ERVK elements with DAXX and aligns with shared ERV repression programs. (**A**) Genomic and repeat annotation of MORC3 ChIP-seq peaks in mESCs. (**B,C**) LTR and ERVK subfamily composition of MORC3 peaks, showing enrichment for ERVK and concentration at IAPEz-int. (**D**) Browser views at representative IAPEz-int loci showing MORC3 signal in wild-type rescue with reduced occupancy in *Daxx* knockout and CSM rescue. (**E**) Structural overlay of MORC2 and MORC3 ATPase domains. (**F**) RNA-seq quantification of DAXX-enriched ERV targets in *Daxx* knockout versus wild-type rescue and in PML or MORC3 perturbations versus matched controls, showing overlap among DAXX-, PML-, and MORC3-dependent ERV silencing programs.

**Figure S8.**
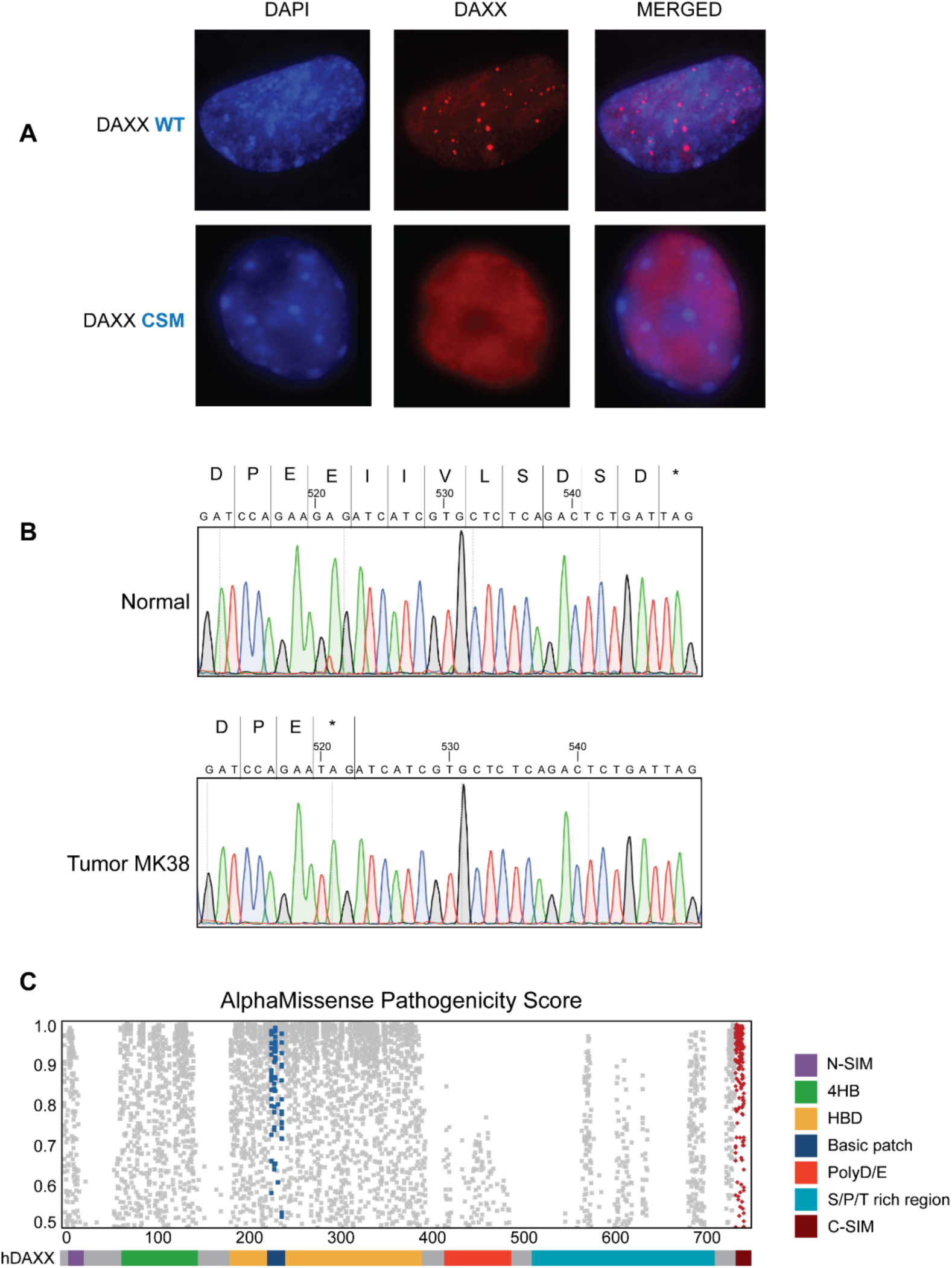
Tumor-linked disruption of the DAXX C-terminal SIM abolishes nuclear puncta and is supported by sequencing and AlphaMissense annotation. (**A**) Immunofluorescence showing punctate nuclear localization of wild-type DAXX and diffuse nuclear signal for the C-SIM mutant (CSM). (**B**) Sanger sequencing of the DAXX C-terminus from normal tissue and tumor MK38, identifying a truncating alteration that deletes the C-SIM. (**C**) AlphaMissense scores across human DAXX, highlighting elevated predicted deleteriousness over the HBD basic surface and the C-terminal SIM tested in this study.

## REFERENCES

1 Allshire, R. C. & Madhani, H. D. Ten principles of heterochromatin formation and function. Nat Rev Mol Cell Biol 19, 229–244 (2018). 10.1038/nrm.2017.119

2 Grewal, S. I. S. The molecular basis of heterochromatin assembly and epigenetic inheritance. Mol Cell 83, 1767–1785 (2023). 10.1016/j.molcel.2023.04.020

3 Padeken, J., Methot, S. P. & Gasser, S. M. Establishment of H3K9-methylated heterochromatin and its functions in tissue differentiation and maintenance. Nat Rev Mol Cell Biol 23, 623–640 (2022). 10.1038/s41580-022-00483-w

4 Du, J., Johnson, L. M., Jacobsen, S. E. & Patel, D. J. DNA methylation pathways and their crosstalk with histone methylation. Nat Rev Mol Cell Biol 16, 519–532 (2015). 10.1038/nrm4043

5 Nicetto, D. et al. H3K9me3-heterochromatin loss at protein-coding genes enables developmental lineage specification. Science 363, 294–297 (2019). 10.1126/science.aau0583

6 Becker, J. S., Nicetto, D. & Zaret, K. S. H3K9me3-Dependent Heterochromatin: Barrier to Cell Fate Changes. Trends Genet 32, 29–41 (2016). 10.1016/j.tig.2015.11.001

7 Stewart-Morgan, K. R., Petryk, N. & Groth, A. Chromatin replication and epigenetic cell memory. Nat Cell Biol 22, 361–371 (2020). 10.1038/s41556-020-0487-y

8 Peng, J. C. & Karpen, G. H. Epigenetic regulation of heterochromatic DNA stability. Curr Opin Genet Dev 18, 204–211 (2008). 10.1016/j.gde.2008.01.021

9 Amaral, N., Ryu, T., Li, X. & Chiolo, I. Nuclear Dynamics of Heterochromatin Repair. Trends Genet 33, 86–100 (2017). 10.1016/j.tig.2016.12.004

10 Groh, S. & Schotta, G. Silencing of endogenous retroviruses by heterochromatin. Cell Mol Life Sci 74, 2055–2065 (2017). 10.1007/s00018-017-2454-8

11 Feschotte, C. Transposable elements and the evolution of regulatory networks. Nat Rev Genet 9, 397–405 (2008). 10.1038/nrg2337

12 Fueyo, R., Judd, J., Feschotte, C. & Wysocka, J. Roles of transposable elements in the regulation of mammalian transcription. Nat Rev Mol Cell Biol 23, 481–497 (2022). 10.1038/s41580-022-00457-y

13 Rowe, H. M. et al. TRIM28 repression of retrotransposon-based enhancers is necessary to preserve transcriptional dynamics in embryonic stem cells. Genome Res 23, 452–461 (2013). 10.1101/gr.147678.112

14 Leung, D. C. & Lorincz, M. C. Silencing of endogenous retroviruses: when and why do histone marks predominate? Trends Biochem Sci 37, 127–133 (2012). 10.1016/j.tibs.2011.11.006

15 Gautam, P., Yu, T. & Loh, Y. H. Regulation of ERVs in pluripotent stem cells and reprogramming. Curr Opin Genet Dev 46, 194–201 (2017). 10.1016/j.gde.2017.07.012

16 Robbez-Masson, L. et al. The HUSH complex cooperates with TRIM28 to repress young retrotransposons and new genes. Genome Res 28, 836–845 (2018). 10.1101/gr.228171.117

17 Muller, I. & Helin, K. Keep quiet: the HUSH complex in transcriptional silencing and disease. Nat Struct Mol Biol 31, 11–22 (2024). 10.1038/s41594-023-01173-7

18 Martire, S. & Banaszynski, L. A. The roles of histone variants in fine-tuning chromatin organization and function. Nat Rev Mol Cell Biol 21, 522–541 (2020). 10.1038/s41580-020-0262-8

19 Geis, F. K. & Goff, S. P. Silencing and Transcriptional Regulation of Endogenous Retroviruses: An Overview. Viruses 12 (2020). 10.3390/v12080884

20 Ahmad, K. & Henikoff, S. The histone variant H3.3 marks active chromatin by replication-independent nucleosome assembly. Mol Cell 9, 1191–1200 (2002). 10.1016/s1097-2765(02)00542-7

21 Park, Y. J. & Luger, K. Histone chaperones in nucleosome eviction and histone exchange. Curr Opin Struct Biol 18, 282–289 (2008). 10.1016/j.sbi.2008.04.003

22 Tyler, J. K. Chromatin assembly. Cooperation between histone chaperones and ATP-dependent nucleosome remodeling machines. Eur J Biochem 269, 2268–2274 (2002). 10.1046/j.1432-1033.2002.02890.x

23 Ray-Gallet, D. et al. Dynamics of histone H3 deposition in vivo reveal a nucleosome gap-filling mechanism for H3.3 to maintain chromatin integrity. Mol Cell 44, 928–941 (2011). 10.1016/j.molcel.2011.12.006

24 Kraushaar, D. C. et al. Genome-wide incorporation dynamics reveal distinct categories of turnover for the histone variant H3.3. Genome Biol 14, R121 (2013). 10.1186/gb-2013-14-10-r121

25 Navarro, C., Lyu, J., Katsori, A. M., Caridha, R. & Elsasser, S. J. An embryonic stem cell-specific heterochromatin state promotes core histone exchange in the absence of DNA accessibility. Nat Commun 11, 5095 (2020). 10.1038/s41467-020-18863-1

26 Dalal, Y., Furuyama, T., Vermaak, D. & Henikoff, S. Structure, dynamics, and evolution of centromeric nucleosomes. Proc Natl Acad Sci U S A 104, 15974–15981 (2007). 10.1073/pnas.0707648104

27 Hammond, C. M., Stromme, C. B., Huang, H., Patel, D. J. & Groth, A. Histone chaperone networks shaping chromatin function. Nat Rev Mol Cell Biol 18, 141–158 (2017). 10.1038/nrm.2016.159

28 Lewis, P. W., Elsaesser, S. J., Noh, K. M., Stadler, S. C. & Allis, C. D. Daxx is an H3.3-specific histone chaperone and cooperates with ATRX in replication-independent chromatin assembly at telomeres. Proc Natl Acad Sci U S A 107, 14075–14080 (2010). 10.1073/pnas.1008850107

29 Drane, P., Ouararhni, K., Depaux, A., Shuaib, M. & Hamiche, A. The death-associated protein DAXX is a novel histone chaperone involved in the replication-independent deposition of H3.3. Genes Dev 24, 1253–1265 (2010). 10.1101/gad.566910

30 Goldberg, A. D. et al. Distinct factors control histone variant H3.3 localization at specific genomic regions. Cell 140, 678–691 (2010). 10.1016/j.cell.2010.01.003

31 Wong, L. H. et al. ATRX interacts with H3.3 in maintaining telomere structural integrity in pluripotent embryonic stem cells. Genome Res 20, 351–360 (2010). 10.1101/gr.101477.109

32 Tagami, H., Ray-Gallet, D., Almouzni, G. & Nakatani, Y. Histone H3.1 and H3.3 complexes mediate nucleosome assembly pathways dependent or independent of DNA synthesis. Cell 116, 51–61 (2004). 10.1016/s0092-8674(03)01064-x

33 Ricketts, M. D. et al. Ubinuclein-1 confers histone H3.3-specific-binding by the HIRA histone chaperone complex. Nat Commun 6, 7711 (2015). 10.1038/ncomms8711

34 Lee, J. S. & Zhang, Z. O-linked N-acetylglucosamine transferase (OGT) interacts with the histone chaperone HIRA complex and regulates nucleosome assembly and cellular senescence. Proc Natl Acad Sci U S A 113, E3213–3220 (2016). 10.1073/pnas.1600509113

35 Elsasser, S. J., Noh, K. M., Diaz, N., Allis, C. D. & Banaszynski, L. A. Histone H3.3 is required for endogenous retroviral element silencing in embryonic stem cells. Nature 522, 240–244 (2015). 10.1038/nature14345

36 He, Q. et al. The Daxx/Atrx Complex Protects Tandem Repetitive Elements during DNA Hypomethylation by Promoting H3K9 Trimethylation. Cell Stem Cell 17, 273–286 (2015). 10.1016/j.stem.2015.07.022

37 Hoelper, D., Huang, H., Jain, A. Y., Patel, D. J. & Lewis, P. W. Structural and mechanistic insights into ATRX-dependent and -independent functions of the histone chaperone DAXX. Nat Commun 8, 1193 (2017). 10.1038/s41467-017-01206-y

38 Sadic, D. et al. Atrx promotes heterochromatin formation at retrotransposons. EMBO Rep 16, 836–850 (2015). 10.15252/embr.201439937

39 Delbarre, E. et al. PML protein organizes heterochromatin domains where it regulates histone H3.3 deposition by ATRX/DAXX. Genome Res 27, 913–921 (2017). 10.1101/gr.215830.116

40 Chang, F. T. et al. PML bodies provide an important platform for the maintenance of telomeric chromatin integrity in embryonic stem cells. Nucleic Acids Res 41, 4447–4458 (2013). 10.1093/nar/gkt114

41 Udugama, M. et al. Histone variant H3.3 provides the heterochromatic H3 lysine 9 tri-methylation mark at telomeres. Nucleic Acids Res 43, 10227–10237 (2015). 10.1093/nar/gkv847

42 Park, J. et al. Long non-coding RNA ChRO1 facilitates ATRX/DAXX-dependent H3.3 deposition for transcription-associated heterochromatin reorganization. Nucleic Acids Res 46, 11759–11775 (2018). 10.1093/nar/gky923

43 Delbarre, E., Ivanauskiene, K., Kuntziger, T. & Collas, P. DAXX-dependent supply of soluble (H3.3-H4) dimers to PML bodies pending deposition into chromatin. Genome Res 23, 440–451 (2013). 10.1101/gr.142703.112

44 Sun, C. et al. The histone chaperone function of Daxx is dispensable for embryonic development. Cell Death Dis 14, 565 (2023). 10.1038/s41419-023-06089-0

45 Wasylishen, A. R. et al. Daxx maintains endogenous retroviral silencing and restricts cellular plasticity in vivo. Sci Adv 6, eaba8415 (2020). 10.1126/sciadv.aba8415

46 Groh, S. et al. Morc3 silences endogenous retroviruses by enabling Daxx-mediated histone H3.3 incorporation. Nat Commun 12, 5996 (2021). 10.1038/s41467-021-26288-7

47 Carraro, M. et al. DAXX adds a de novo H3.3K9me3 deposition pathway to the histone chaperone network. Mol Cell 83, 1075–1092 e1079 (2023). 10.1016/j.molcel.2023.02.009

48 Jiao, Y. et al. DAXX/ATRX, MEN1, and mTOR pathway genes are frequently altered in pancreatic neuroendocrine tumors. Science 331, 1199–1203 (2011). 10.1126/science.1200609

49 Heaphy, C. M. et al. Altered telomeres in tumors with ATRX and DAXX mutations. Science 333, 425 (2011). 10.1126/science.1207313

50 Dyer, M. A., Qadeer, Z. A., Valle-Garcia, D. & Bernstein, E. ATRX and DAXX: Mechanisms and Mutations. Cold Spring Harb Perspect Med 7 (2017). 10.1101/cshperspect.a026567

51 de Wilde, R. F. et al. Loss of ATRX or DAXX expression and concomitant acquisition of the alternative lengthening of telomeres phenotype are late events in a small subset of MEN-1 syndrome pancreatic neuroendocrine tumors. Mod Pathol 25, 1033–1039 (2012). 10.1038/modpathol.2012.53

52 Marinoni, I. et al. Loss of DAXX and ATRX are associated with chromosome instability and reduced survival of patients with pancreatic neuroendocrine tumors. Gastroenterology 146, 453–460 e455 (2014). 10.1053/j.gastro.2013.10.020

53 Clynes, D. et al. Suppression of the alternative lengthening of telomere pathway by the chromatin remodelling factor ATRX. Nat Commun 6, 7538 (2015). 10.1038/ncomms8538

54 Singhi, A. D. et al. Alternative Lengthening of Telomeres and Loss of DAXX/ATRX Expression Predicts Metastatic Disease and Poor Survival in Patients with Pancreatic Neuroendocrine Tumors. Clin Cancer Res 23, 600–609 (2017). 10.1158/1078-0432.CCR-16-1113

55 Gao, J., Liu, T., Yang, D. & Tuo, Q. The Dynamic Regulation of Daxx-Mediated Transcriptional Inhibition by SUMO and PML NBs. Int J Mol Sci 26 (2025). 10.3390/ijms26146703

56 Rowe, H. M. et al. KAP1 controls endogenous retroviruses in embryonic stem cells. Nature 463, 237–240 (2010). 10.1038/nature08674

57 Matsui, T. et al. Proviral silencing in embryonic stem cells requires the histone methyltransferase ESET. Nature 464, 927–931 (2010). 10.1038/nature08858

58 Sachs, P. et al. SMARCAD1 ATPase activity is required to silence endogenous retroviruses in embryonic stem cells. Nat Commun 10, 1335 (2019). 10.1038/s41467-019-09078-0

59 Schotta, G. et al. A silencing pathway to induce H3-K9 and H4-K20 trimethylation at constitutive heterochromatin. Genes Dev 18, 1251–1262 (2004). 10.1101/gad.300704

60 Elsasser, S. J. et al. DAXX envelops a histone H3.3-H4 dimer for H3.3-specific recognition. Nature 491, 560–565 (2012). 10.1038/nature11608

61 Liu, C. P. et al. Structure of the variant histone H3.3-H4 heterodimer in complex with its chaperone DAXX. Nat Struct Mol Biol 19, 1287–1292 (2012). 10.1038/nsmb.2439

62 DeNizio, J. E., Elsasser, S. J. & Black, B. E. DAXX co-folds with H3.3/H4 using high local stability conferred by the H3.3 variant recognition residues. Nucleic Acids Res 42, 4318–4331 (2014). 10.1093/nar/gku090

63 Zink, L. M. et al. H3.Y discriminates between HIRA and DAXX chaperone complexes and reveals unexpected insights into human DAXX-H3.3-H4 binding and deposition requirements. Nucleic Acids Res 45, 5691–5706 (2017). 10.1093/nar/gkx131

64 Sauer, P. V. et al. Insights into the molecular architecture and histone H3-H4 deposition mechanism of yeast Chromatin assembly factor 1. Elife 6 (2017). 10.7554/eLife.23474

65 Liu, C. P. et al. Structural insights into histone binding and nucleosome assembly by chromatin assembly factor-1. Science 381, eadd8673 (2023). 10.1126/science.add8673

66 Mattiroli, F. et al. DNA-mediated association of two histone-bound complexes of yeast Chromatin Assembly Factor-1 (CAF-1) drives tetrasome assembly in the wake of DNA replication. Elife 6 (2017). 10.7554/eLife.22799

67 Rosas, R. et al. A novel single alpha-helix DNA-binding domain in CAF-1 promotes gene silencing and DNA damage survival through tetrasome-length DNA selectivity and spacer function. Elife 12 (2023). 10.7554/eLife.83538

68 Liu, W. H., Roemer, S. C., Port, A. M. & Churchill, M. E. CAF-1-induced oligomerization of histones H3/H4 and mutually exclusive interactions with Asf1 guide H3/H4 transitions among histone chaperones and DNA. Nucleic Acids Res 40, 11229–11239 (2012). 10.1093/nar/gks906

69 Rouillon, C. et al. CAF-1 deposits newly synthesized histones during DNA replication using distinct mechanisms on the leading and lagging strands. Nucleic Acids Res 51, 3770–3792 (2023). 10.1093/nar/gkad171

70 Cheng, J. et al. Accurate proteome-wide missense variant effect prediction with AlphaMissense. Science 381, eadg7492 (2023). 10.1126/science.adg7492

71 Chan, C. S. et al. ATRX, DAXX or MEN1 mutant pancreatic neuroendocrine tumors are a distinct alpha-cell signature subgroup. Nat Commun 9, 4158 (2018). 10.1038/s41467-018-06498-2

72 Huang, L. et al. DAXX represents a new type of protein-folding enabler. Nature 597, 132–137 (2021). 10.1038/s41586-021-03824-5

73 Mukhopadhyay, D. & Matunis, M. J. SUMmOning Daxx-mediated repression. Mol Cell 42, 4–5 (2011). 10.1016/j.molcel.2011.03.008

74 Sloan, E., Orr, A. & Everett, R. D. MORC3, a Component of PML Nuclear Bodies, Has a Role in Restricting Herpes Simplex Virus 1 and Human Cytomegalovirus. J Virol 90, 8621–8633 (2016). 10.1128/JVI.00621-16

75 Desai, V. P. et al. The role of MORC3 in silencing transposable elements in mouse embryonic stem cells. Epigenetics Chromatin 14, 49 (2021). 10.1186/s13072-021-00420-9

76 Andrews, A. J., Chen, X., Zevin, A., Stargell, L. A. & Luger, K. The histone chaperone Nap1 promotes nucleosome assembly by eliminating nonnucleosomal histone DNA interactions. Mol Cell 37, 834–842 (2010). 10.1016/j.molcel.2010.01.037

77 D’Arcy, S. et al. Chaperone Nap1 shields histone surfaces used in a nucleosome and can put H2A-H2B in an unconventional tetrameric form. Mol Cell 51, 662–677 (2013). 10.1016/j.molcel.2013.07.015

78 Zhang, K. et al. A DNA binding winged helix domain in CAF-1 functions with PCNA to stabilize CAF-1 at replication forks. Nucleic Acids Res 44, 5083–5094 (2016). 10.1093/nar/gkw106

79 Mattiroli, F., Gu, Y., Balsbaugh, J. L., Ahn, N. G. & Luger, K. The Cac2 subunit is essential for productive histone binding and nucleosome assembly in CAF-1. Sci Rep 7, 46274 (2017). 10.1038/srep46274

80 Liu, Y. et al. Structural analysis of Rtt106p reveals a DNA binding role required for heterochromatin silencing. J Biol Chem 285, 4251–4262 (2010). 10.1074/jbc.M109.055996

81 Huang, S. et al. Rtt106p is a histone chaperone involved in heterochromatin-mediated silencing. Proc Natl Acad Sci U S A 102, 13410–13415 (2005). 10.1073/pnas.0506176102

82 Imbeault, D., Gamar, L., Rufiange, A., Paquet, E. & Nourani, A. The Rtt106 histone chaperone is functionally linked to transcription elongation and is involved in the regulation of spurious transcription from cryptic promoters in yeast. J Biol Chem 283, 27350–27354 (2008). 10.1074/jbc.C800147200

83 Zunder, R. M., Antczak, A. J., Berger, J. M. & Rine, J. Two surfaces on the histone chaperone Rtt106 mediate histone binding, replication, and silencing. Proc Natl Acad Sci U S A 109, E144–153 (2012). 10.1073/pnas.1119095109

84 Fazly, A. et al. Histone chaperone Rtt106 promotes nucleosome formation using (H3-H4)2 tetramers. J Biol Chem 287, 10753–10760 (2012). 10.1074/jbc.M112.347450

85 Ricketts, M. D. et al. The HIRA histone chaperone complex subunit UBN1 harbors H3/H4- and DNA-binding activity. J Biol Chem 294, 9239–9259 (2019). 10.1074/jbc.RA119.007480

86 Ray-Gallet, D. et al. Functional activity of the H3.3 histone chaperone complex HIRA requires trimerization of the HIRA subunit. Nat Commun 9, 3103 (2018). 10.1038/s41467-018-05581-y

87 Torne, J. et al. Two HIRA-dependent pathways mediate H3.3 de novo deposition and recycling during transcription. Nat Struct Mol Biol 27, 1057–1068 (2020). 10.1038/s41594-020-0492-7

88 Pchelintsev, N. A. et al. Placing the HIRA histone chaperone complex in the chromatin landscape. Cell Rep 3, 1012–1019 (2013). 10.1016/j.celrep.2013.03.026

89 Wolf, D. & Goff, S. P. TRIM28 mediates primer binding site-targeted silencing of murine leukemia virus in embryonic cells. Cell 131, 46–57 (2007). 10.1016/j.cell.2007.07.026

90 Ecco, G. et al. Transposable Elements and Their KRAB-ZFP Controllers Regulate Gene Expression in Adult Tissues. Dev Cell 36, 611–623 (2016). 10.1016/j.devcel.2016.02.024

91 Imbeault, M., Helleboid, P. Y. & Trono, D. KRAB zinc-finger proteins contribute to the evolution of gene regulatory networks. Nature 543, 550–554 (2017). 10.1038/nature21683

92 Voon, H. P. et al. ATRX Plays a Key Role in Maintaining Silencing at Interstitial Heterochromatic Loci and Imprinted Genes. Cell Rep 11, 405–418 (2015). 10.1016/j.celrep.2015.03.036

93 Fang, Y. et al. ATRX guards against aberrant differentiation in mesenchymal progenitor cells. Nucleic Acids Res 52, 4950–4968 (2024). 10.1093/nar/gkae160

94 Iwase, S. et al. ATRX ADD domain links an atypical histone methylation recognition mechanism to human mental-retardation syndrome. Nat Struct Mol Biol 18, 769–776 (2011). 10.1038/nsmb.2062

95 Dhayalan, A. et al. The ATRX-ADD domain binds to H3 tail peptides and reads the combined methylation state of K4 and K9. Hum Mol Genet 20, 2195–2203 (2011). 10.1093/hmg/ddr107

96 Wang, X., Zhao, Y., Zhang, J. & Chen, Y. Structural basis for DAXX interaction with ATRX. Protein Cell 8, 767–771 (2017). 10.1007/s13238-017-0462-y

97 Lin, D. Y. et al. Role of SUMO-interacting motif in Daxx SUMO modification, subnuclear localization, and repression of sumoylated transcription factors. Mol Cell 24, 341–354 (2006). 10.1016/j.molcel.2006.10.019

98 Escobar-Cabrera, E. et al. Characterizing the N- and C-terminal Small ubiquitin-like modifier (SUMO)-interacting motifs of the scaffold protein DAXX. J Biol Chem 286, 19816–19829 (2011). 10.1074/jbc.M111.231647

99 Shastrula, P. K. et al. PML is recruited to heterochromatin during S phase and represses DAXX-mediated histone H3.3 chromatin assembly. J Cell Sci 132 (2019). 10.1242/jcs.220970

100 Chang, C. C. et al. Structural and functional roles of Daxx SIM phosphorylation in SUMO paralog-selective binding and apoptosis modulation. Mol Cell 42, 62–74 (2011). 10.1016/j.molcel.2011.02.022

101 Ivanov, A. V. et al. PHD domain-mediated E3 ligase activity directs intramolecular sumoylation of an adjacent bromodomain required for gene silencing. Mol Cell 28, 823–837 (2007). 10.1016/j.molcel.2007.11.012

102 Li, X. et al. SUMOylation of the transcriptional co-repressor KAP1 is regulated by the serine and threonine phosphatase PP1. Sci Signal 3, ra32 (2010). 10.1126/scisignal.2000781

103 Schultz, D. C., Ayyanathan, K., Negorev, D., Maul, G. G. & Rauscher, F. J., 3rd. SETDB1: a novel KAP-1-associated histone H3, lysine 9-specific methyltransferase that contributes to HP1-mediated silencing of euchromatic genes by KRAB zinc-finger proteins. Genes Dev 16, 919–932 (2002). 10.1101/gad.973302

104 Tan, W. et al. MORC2 is a phosphorylation-dependent DNA compaction machine. Nat Commun 16, 5606 (2025). 10.1038/s41467-025-60751-z

105 Kim, H. et al. The Gene-Silencing Protein MORC-1 Topologically Entraps DNA and Forms Multimeric Assemblies to Cause DNA Compaction. Mol Cell 75, 700–710 e706 (2019). 10.1016/j.molcel.2019.07.032

106 Hertwig, F., Peifer, M. & Fischer, M. Telomere maintenance is pivotal for high-risk neuroblastoma. Cell Cycle 15, 311–312 (2016). 10.1080/15384101.2015.1125243

107 Cai, J. et al. ATRX mRNA expression combined with IDH1/2 mutational status and Ki-67 expression refines the molecular classification of astrocytic tumors: evidence from the whole transcriptome sequencing of 169 samples samples. Oncotarget 5, 2551–2561 (2014). 10.18632/oncotarget.1838

108 Mason-Osann, E. et al. Identification of a novel gene fusion in ALT positive osteosarcoma. Oncotarget 9, 32868–32880 (2018). 10.18632/oncotarget.26029

109 Yang, C. Y. et al. Targeted next-generation sequencing of cancer genes identified frequent TP53 and ATRX mutations in leiomyosarcoma. Am J Transl Res 7, 2072–2081 (2015).

110 Slatter, T. L. et al. Loss of ATRX and DAXX expression identifies poor prognosis for smooth muscle tumours of uncertain malignant potential and early stage uterine leiomyosarcoma. J Pathol Clin Res 1, 95–105 (2015). 10.1002/cjp2.11

111 Clatterbuck Soper, S. F., et al. Cancer-associated DAXX mutations reveal a critical role for ATRX localization in ALT suppression. bioRxiv (2024). 10.1101/2024.11.18.624165

112 Michaelson, J. S., Bader, D., Kuo, F., Kozak, C. & Leder, P. Loss of Daxx, a promiscuously interacting protein, results in extensive apoptosis in early mouse development. Genes Dev 13, 1918–1923 (1999). 10.1101/gad.13.15.1918

113 Langmead, B., Trapnell, C., Pop, M. & Salzberg, S. L. Ultrafast and memory-efficient alignment of short DNA sequences to the human genome. Genome Biol 10, R25 (2009). 10.1186/gb-2009-10-3-r25

114 Li, H. et al. The Sequence Alignment/Map format and SAMtools. Bioinformatics 25, 2078–2079 (2009). 10.1093/bioinformatics/btp352

115 Ramirez, F., Dundar, F., Diehl, S., Gruning, B. A. & Manke, T. deepTools: a flexible platform for exploring deep-sequencing data. Nucleic Acids Res 42, W187–191 (2014). 10.1093/nar/gku365

116 Feng, J., Liu, T., Qin, B., Zhang, Y. & Liu, X. S. Identifying ChIP-seq enrichment using MACS. Nat Protoc 7, 1728–1740 (2012). 10.1038/nprot.2012.101

117 Heinz, S. et al. Simple combinations of lineage-determining transcription factors prime cis-regulatory elements required for macrophage and B cell identities. Mol Cell 38, 576–589 (2010). 10.1016/j.molcel.2010.05.004

118 Thorvaldsdottir, H., Robinson, J. T. & Mesirov, J. P. Integrative Genomics Viewer (IGV): high-performance genomics data visualization and exploration. Brief Bioinform 14, 178–192 (2013). 10.1093/bib/bbs017

119 Kawase, M. & Ichiyanagi, K. Mouse retrotransposons: sequence structure, evolutionary age, genomic distribution and function. Genes Genet Syst 98, 337–351 (2024). 10.1266/ggs.23-00221

120 Brattas, P. L. et al. TRIM28 Controls a Gene Regulatory Network Based on Endogenous Retroviruses in Human Neural Progenitor Cells. Cell Rep 18, 1–11 (2017). 10.1016/j.celrep.2016.12.010

121 Dobin, A. et al. STAR: ultrafast universal RNA-seq aligner. Bioinformatics 29, 15–21 (2013). 10.1093/bioinformatics/bts635

122 Jin, Y., Tam, O. H., Paniagua, E. & Hammell, M. TEtranscripts: a package for including transposable elements in differential expression analysis of RNA-seq datasets. Bioinformatics 31, 3593–3599 (2015). 10.1093/bioinformatics/btv422

123 Love, M. I., Huber, W. & Anders, S. Moderated estimation of fold change and dispersion for RNA-seq data with DESeq2. Genome Biol 15, 550 (2014). 10.1186/s13059-014-0550-8

124 Ito, K. & Murphy, D. Application of ggplot2 to Pharmacometric Graphics. CPT Pharmacometrics Syst Pharmacol 2, e79 (2013). 10.1038/psp.2013.56

125 Quinlan, A. R. & Hall, I. M. BEDTools: a flexible suite of utilities for comparing genomic features. Bioinformatics 26, 841–842 (2010). 10.1093/bioinformatics/btq033

126 Jumper, J. et al. Highly accurate protein structure prediction with AlphaFold. Nature 596, 583–589 (2021). 10.1038/s41586-021-03819-2

127 Hornbeck, P. V. et al. PhosphoSitePlus, 2014: mutations, PTMs and recalibrations. Nucleic Acids Res 43, D512–520 (2015). 10.1093/nar/gku1267

